# Proteogenomic characterization of age and sex interactions in cardiac gene expression

**DOI:** 10.1101/2022.05.11.491401

**Authors:** Yu Han, Sara A. Wennersten, Julianna Wright, R. W. Ludwig, Edward Lau, Maggie P. Y. Lam

## Abstract

The risks of heart diseases are significantly modulated by biological age and sex, but how these factors influence baseline cardiac gene expression remains incompletely understood. Here we characterized young adult and early aging mouse hearts using proteogenomics to identify age and sex dependent gene expression signatures in the mouse heart. RNA sequencing from 4 months old and 20 months old female and male C57BL/6J hearts identified thousands of genes with differential transcript abundances both between sexes (male vs. female) and across age groups (20 mo. vs. 4 mo.). Sex-associated cardiac genes are broadly distributed, functioning in the TCA cycle, mitochondrial translation, autophagy, and other processes. In addition, we found over 800 genes with differential aging response between male and female, which are enriched in calmodulin signaling and cell cycle regulations. Comparison with mass spectrometry data shows a cluster of metabolism genes with up-regulated transcripts but down-regulated protein levels in aging hearts, consistent with an uncoupling of transcriptional regulations in the genetic program with protein compositions. An analysis of sex-adjusted aging cardiac transcriptomes further revealed widespread remodeling of exon usage patterns that is largely independent from differential gene expression, concomitant with upstream changes in RNA-binding protein and splice factor transcripts. To evaluate the potential impact of the splicing events on proteoform composition in the heart, we applied an RNA-guided-proteomics computational pipeline to analyze the mass spectrometry data, and putatively identified hundreds of splice events with the potential to rewire the cardiac proteome through producing detectable splice isoform specific peptides. Taken together, this study contributes to emerging evidences for considerable sexual dimorphism in the cardiac aging process involving sex-biased aging genes and regulatory networks. Aging hearts are associated with a rewiring of RNA splicing programs, including sex- and age-dependent changes in exon usages and splice patterns that have the potential to influence cardiac protein structure and function. These changes represent an under-investigated aspect of cardiac aging that should be considered in the search for disease mechanisms.

## Introduction

There are considerable sex differences in the risk, prevalence, and prognosis of heart diseases (Leinwand, 2003). The effect of sex is age dependent; whereas pre-menopausal adult women are relatively protected from cardiovascular diseases compared to age-matched men, older women at post-menopausal ages experience higher risks for adverse cardiac remodeling and higher incidences of heart failure with preserved ejection fraction (Barton and Meyer, 2020; Cheng et al., 2010; Sotomi et al., 2021). Recent work has centered on considerable sexually dimorphisms in aging between men and women that underlie clinical observations, where biological sex exerts modulating influence on the fundamental biology of cardiac aging processes (Ji et al., 2022). Among other molecular and phenotypic differences, aging women have higher increase in left ventricular wall thickness and concentric remodeling and than men (Cheng et al., 2010; Gebhard et al., 2013), increased collagen deposition, and experience more diastolic dysfunction (Ji et al., 2022; Sotomi et al., 2021). The limited efficacy of hormone replacement therapy in remediating sex-specific risks is consistent with multifactorial underlying causes that should be considered based on intrinsic differences of female and male hearts. Targeting cardiac aging-related gene regulatory networks has been proposed as a promising therapeutic approach against heart diseases (Chen et al., 2022; Ji et al., 2022), but progress will likely require more comprehensive understanding of the interacting effects of both biological age and sex on cardiac gene expression.

In rodents, female and male hearts show intrinsic differences in cell proportion, cell morphology, and myocyte contractility (Squiers et al., 2021; Walker et al., 2021). Several hundreds of cardiac genes have previously been found to display sexual dimorphism in expression levels (Trexler et al., 2017). Differences between male and female hearts may be attributable to gonadal hormonal differences as well as developmental differences driven by sex chromosomes (DeLeon-Pennell and Lindsey, 2019; Shi et al., 2021; Squiers et al., 2021). Paralleling humans, young female mice are generally protected and have better survivals in multiple cardiac remodeling and failure models, whereas older females experience steeper functional decline during cardiac aging (DuPont et al., 2021). Here, we investigated the effect of age and sex associations on gene expression in the mouse heart, by performing a proteogenomic characterization of 4 mo. vs. 20 mo. old female and male C57BL/6J mice, with the latter age group corresponding roughly to the seventh decade of life in human age (Flurkey et al., 2007). With this study design, we compared baseline sex differences and sex-adjusted effects of aging as well as genes showing age-sex interactions. The results indicate that both sex and age exert significant effect on the expression levels of thousands of cardiac genes, and moreover there are significant sex-specific effects on age-associated changes in gene expression involving calmodulin signaling and cell cycle regulation pathways. Latent space representation of the sex-adjusted age comparisons revealed gene regulatory modules in cardiac aging involving RNA-binding proteins, concomitant with a widespread remodeling of exon usage patterns, including splice events that can be linked to proteoform composition at the proteome level. These findings expand the current understanding on the molecular underpinnings of sex differences in cardiac phenotypes and portrait a complex proteogenomic landscape of sex-dependent changes in chronological aging in mice.

## Methods

### Animal Models

All animal protocols were approved by the Institutional Animal Care and Use Committee (IACUC) guidelines at the University of Colorado Denver/Anschutz Medical Campus. Wild-type C57BL/6J mice were purchased from Jackson Laboratories (Bar Harbor, ME, USA) and housed in a temperature-controlled environment on a 12-hr light/dark cycle and fed with normal diet and water ad libitum under National Institutes of Health (NIH) guidelines for the Care and Use of Laboratory Animals. Young adult (4 mo.) vs. early aging (20 mo.) mice (male and female, n = 5) were examined by echocardiographic analysis and sacrificed. Body weight, heart weight and tibia length were measured. Heart ventricles were collected and kept frozen at –80 °C.

### Echocardiographic analysis

Transthoracic echocardiograms were performed using the VisualSonics Vevo2100 system (Visualsonics, Toronto, Canada). Animals were anesthetized in an induction chamber at 2% isoflurane. Once appropriately sedated, mice were transferred to the imaging platform. Hair on the chest was removed with a depilatory lotion and the mouse was secured by taping all four limbs to the platform. Body temperature was maintained at 37°C. Body temperature of the mouse was monitored through a rectal probe connected to the Advanced Physiological Monitoring Unit, which also displays and records the animal’s ECG and Respiration Rate. Isoflurane was delivered through a nose cone at 1.5% throughout the imaging session. Mice were under anesthesia for no longer than 15 minutes. Parasternal long axis (PSLAX) and parasternal short axis (SAX) views of the left ventricle (LV) were acquired using B-mode imaging. SAX views of the LV at the papillary muscle level were used to acquire M-mode images. Left ventricular anterior wall (LVAW) thickness, left ventricular posterior wall (LVPW) thickness, and left ventricular internal diameter (LVID) in systole and diastole were measured using the M-mode images in order to measure changes in wall thickness and systolic function (**Supplemental Table S1**). Heart rate was measured in each M-mode image and averaged. All measurements were averaged from at least three cardiac cycles during the exhale phase and taken at body temperatures between 37°C and 38°C. All echocardiographic measurements and analyses were performed in a blinded manner.

### RNA extraction and sequencing

To extract total RNA, frozen tissues were weighed, thawed, and excised into small cubes (2–3 mm^3^) on weighing boats on ice. Tissue pieces were transferred to 2.0 mL RNase-free eppendorf tubes. Cold TRIzol (Invitrogen catalog #15596026) was added to tissues at a ratio of 75:1 (75 μL TRIzol for 1 mg of tissue). Samples were homogenized with a handheld homogenizer (OMNI International TH) for 10 seconds, followed by 1 minute cooling on ice, repeated for a total of 3 rounds of homogenization. The samples were centrifuged at 14,000 ×g for 15 min at 4°C after which supernatants were transferred to a new tube and proceed to RNA extraction using Direct-zol RNA kit (ZYMO #R2072) following manufacturer’s instructions. RNA concentration and quality were measured on a Nanodrop instrument (Thermo #ND-2000). PolyA+ mRNA were selected using oligo-dT beads and libraries were constructed with NEBNext Ultra II RNA Library Prep (unstranded) (Illumina). Paired-end sequencing (150 bp reads) was performed on a NovaSeq 6000 Sequencing System (Illumina). Untrimmed raw reads (fastq, 150 nt PE) were mapped using STAR v.2.7.6a (Dobin et al., 2013) against the GRCm39 genome assembly and GENCODE M26 (Frankish et al., 2021) annotations. Aligned reads were assembled into transcripts using Stringtie v.2.1.1 to output the gene and transcript count matrices. From the gene count matrix files, we performed differential gene expression analysis using DESeq2 v.1.30.1 (Love et al., 2014) in R v.4.0.4 with the formula “∽sex + age + age:sex”, followed by fold-change shrinkage using apeglm (Zhu et al., 2019) to estimate logFC and false discovery rate (FDR)-adjusted s-value (false sign or small) for |logFC| ≥ 0.1.

### Protein extraction and mass spectrometry

Frozen tissues were weighed, thawed, and cut into small pieces (2–3 mm^3^) on weigh boats on ice. Tissue pieces were transferred to 2.0 mL eppendorf tubes. Cold RIPA lysis buffer (Thermo #89901) with protease inhibitors (Pierce Halt) was added to tissues at a ratio of 20:1 (20 μL RIPA per 1 mg of tissue) and samples were homogenized as above then sonicated with a handheld sonicator (Fisher #FB120110) at 40% amplitude for 15 cycles (each cycle is 1 s of sonication, followed by 5 s pause; three rounds. The samples were centrifuged at 14,000 × g for 15 min at 4°C, and the supernatants were collected. Soluble protein concentration was measured with the BCA protein assay kit following manufacturer instruction; 100 μg of proteins were digested using a filter-assisted protocol as described (Manza et al., 2005) with a 3-hour LysC digestion and 12-hr trypsin digestion. Peptide digest were tagged with tandem mass tags (TMT; 10-plex reagent, Thermo #90110) following manufacturer’s instructions, using a random number generator to assign balanced randomized samples across TMT blocks and channels. Tagged digests within the same block are then mixed and fractionated with the high-pH reversed phase peptide fractionation kit (Thermo #84868). Fractionated peptides are dried and resuspended in 0.1% formic acid (LC-MS grade, thermo #28905) at a final concentration of 0.5 μg/μL.

Each peptide fraction (∽1.5 μg) was further separated with online low-pH reversed-phase LC (PepMap C18 column, 3-μm particle, 100-Å pore; 75 μm x 150 mm; Thermo Fisher Scientific) via the EASYnLC 1200 system coupled to the Easy-Spray ion source (Thermo Fisher Scientific) at 300 nL/min with a 120 min gradient: 0 -105 min: 0 to 40% B; 105 - 110 min: 40 to 70% B; 110 - 115 min: 70 to 100% B; 115 - 120 min: 100% B (solvent A: 0.1% v/v formic acid; solvent B: 80% v/v acetonitrile; column temperature: 50 °C). Mass spectra were acquired on a Thermo Scientific Q-Exactive HF Orbitrap mass spectrometer with the following settings: polarity: positive; data dependent acquisition (DDA): top 15 ions, MS resolution: 60,000, mass range: 300-1650 m/z; precursor dynamic exclusion: 30s, maximum ion injection time: 20 ms; MS automatic gain control (AGC) target: 3e6. isolation window: 1.4 m/z; stepped normalized collision energy (NCE): 28, 30, 32. MS2 resolution; 60,000; MS2 maximum ion injection time: 100 ms; MS2 AGC target: 2e5. Raw mass spectrometry data were converted to mzML format using ThermoRawFileParser v.1.2.0 (Hulstaert et al., 2020) then searched against UniProt SwissProt (UniProt Consortium, 2021) *Mus musculus* sequences (v2021_02) using Comet v.2020_013 (Eng et al., 2015). Database search results were post-processed using Percolator v.3 (Crux v.4.0 distribution) (The et al., 2016) requiring 1% FDR for identification. TMT intensities were extracted from MS2 spectra using a custom Python script (Dostal et al., 2020) built on the pymzml library (Kösters et al., 2018) then corrected for isotope contamination and normalized across blocks as previously described (Han et al., 2021).

### Functional enrichment and pathway analysis

Pathway enrichment was performed using Fisher’s exact test against Reactome terms using ReactomePA (Yu and He, 2016) and GSEA using the fgsea package (Korotkevich et al., 2016) against MSigDB Canonical pathways (Liberzon et al., 2015). For GSVA, for each gene set, we used the gsva package to calculate an enrichment score in each sample, which was then used to perform differential enrichment analysis by limma. An integer normalized count matrix was created from the DESeq2 data, after which mouse Ensembl was queried with biomaRt (Durinck et al., 2009) to retrieve the gene name from the Ensembl Gene ID. We treated human and mouse genes sharing identical gene names as orthologs, and used the gene names to retrieve human Entrez Gene ID from the human Ensembl Biomart. We then used the Broad MSigDB ontology gene set collection v7.4 to construct the GSVA dataset. For PLIER, starting with the DESeq2 dataset again, we retrieved the VST transformed data or the normalized counts. Because of the small sample size, we used a transfer learning framework to generalize the PLIER method by allowing the pathway loading matrix to be first trained in a large collection of RNA-seq data containing gene expression latent structural information then transferred to the collected data set (Taroni et al., 2019). To do so, we used GTEx v8 (The GTEx Consortium, 2020) RNA sequencing data (∽17,000 samples) to learn the latent variable-pathway mapping and train a PLIER model, employing the built-in regularizations to favor latent variables that map to only a few human interpretable pathways (Mao et al., 2019). Differential regulation of the gene sets and latent variables were then assessed using limma v.3.50.1 (Ritchie et al., 2015) with age and sex as factors in the model.

### Exon usage and RNA-guided protein sequence databases

Exon usage analysis was performed with the aid of DEXSeq v.1.40 (Anders et al., 2012). The flattened annotation file was generated with the DEXSeq Python scripts using GENCODE M26 gtf annotation. Differential exon usage was estimated using a ∽sample + exon + age:exon + sex:exon + age:sex:exon design.

To generate custom protein sequence databases containing sample-specific splice junctions, we used a Python program we developed (JCAST.py) (Lau et al., 2019; Ludwig and Lau, 2021). Briefly, we used rMATS v.4.1.0 (Shen et al., 2014) with --read-length 150 option and the GENCODE M26 gtf annotation to model alternative splicing events in all gathered RNA sequencing data. From the resulting 31,899 splice junctions belonging to 8712 genes, we performed one-frame in silico translation using Ensembl annotated translation start sites and frames to produce custom protein sequences corresponding to likely translatable splice junctions. In total, we translated 6,295 non-canonical sequences that did not experience frameshift (jcast_t1) and can be joined from the upstream and downstream exons back to the canonical Uniprot sequence via 10-residue flanks, among which 5412 are not found in the mouse SwissProt canonical + isoform databases. Translations that experience frameshift regardless of whether they encounter premature termination codons (jcast_t2–t4) and those that cannot be merged back to SwissProt canonical sequences (orphans) were discarded (Han et al., 2020; Lau et al., 2019). We then appended the translated non-canonical jcast_t1 sequences to canonical mouse entries and searched the labeled tandem mass spectrometry experiments on the mouse hearts.

### Structure prediction and additional analyses

To predict isoform structure, folded 3D protein models were generated from a .fasta file with the “AlphaFold2_advanced” Google Colab notebook (Jumper et al., 2021; Mirdita et al., 2021). The resulting highest-ranked .pdb file was used as input to the DeepFRI web server (Gligorijevic et al., 2019) to predict protein function. The functional salience map was used to estimate functional importance of individual residues. Additional statistical analysis and visualization was performed in R v.4.1.3 with the aid of the ggpubr (Kassambara, 2021) and ggsankey packages.

### Data availability

RNA sequencing data have been deposited to NCBI GEO. Mass spectrometry data have been deposited to ProteomeXchange (Deutsch et al., 2020) via jPOST (Okuda et al., 2017).

## Results and Discussions

### Sex and age associated differential gene expression and in the mouse heart

We performed echocardiographic and proteogenomics of male and female 20 mo. vs. 4 mo. C57BL6/J mice (n = 3–5 per group) (**Figure 1a**). The age groups are consistent with previous aging studies in the mouse, with the latter thought to correspond to the seventh decade of human life based on cumulative mortality (Flurkey et al., 2007). In our hands, aged animals at 20 months of age showed an increase in left ventricular mass over tibia length as well as systolic and diastolic left ventricular anterior wall thickness (**Figure 1b**), consistent with age-associated left ventricular hypertrophy. Echocardiography shows no significant difference in systolic function in the animals but a difference in diastolic function especially in male animals (**Figure 1c**). To compare age and sex differences in the transcriptomic, we performed polyA+ RNA sequencing from 12 of the mouse heart samples (3 male and 3 female at 4 months; 3 male and 3 female at 20 months).

**Figure 1:**
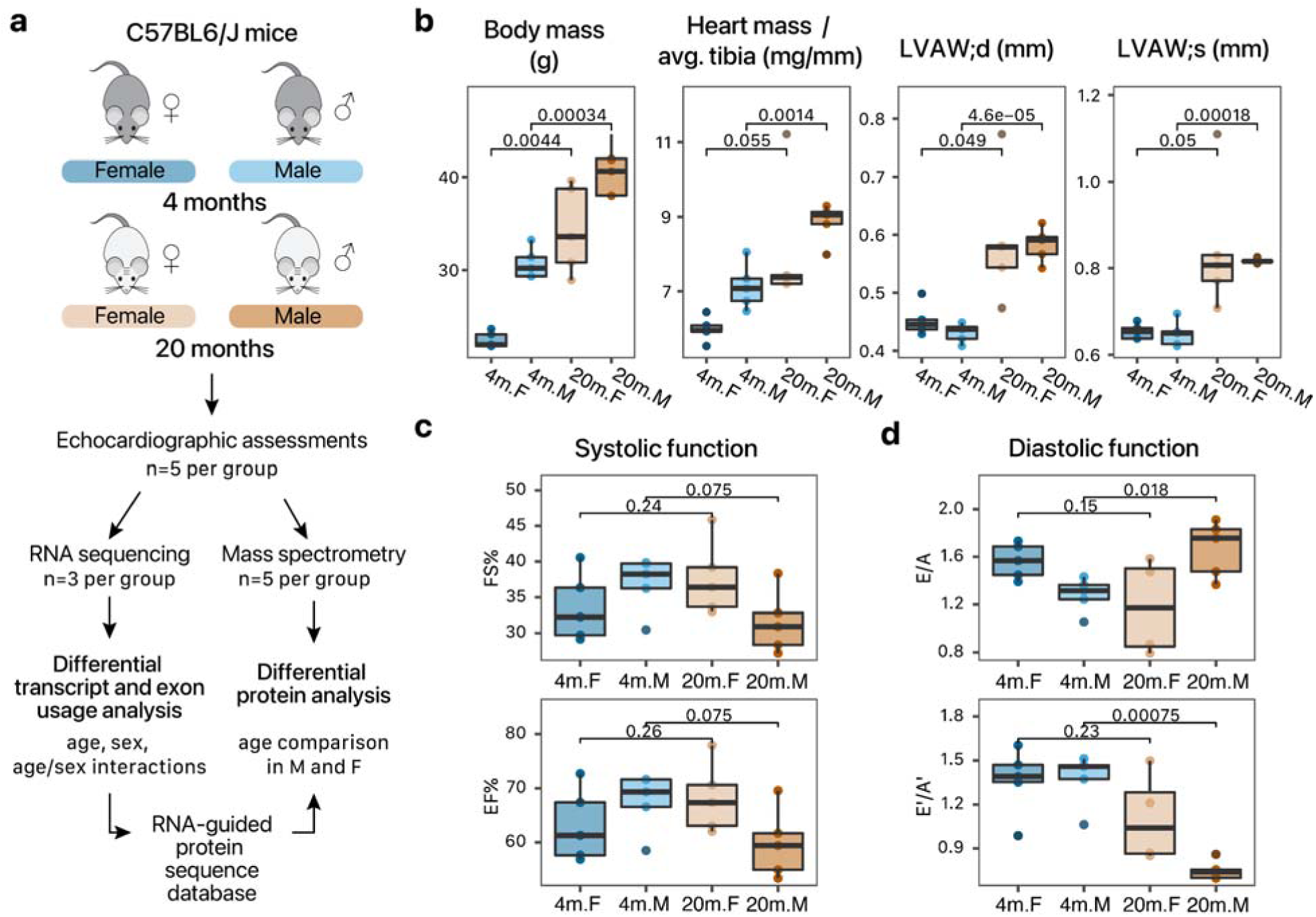
Study design. **a**. Animal groups and schema of proteogenomics workflow. **b**. Body mass, heart mass, anterior wall thicknesses in each group (n=5 biological replicates per group). P values: Student’s t test. **c–d**. Echocardiographic measures associated with c. systolic and d. diastolic function.

The first two principal components of the RNA sequencing data showed clear separation of transcriptome profiles by sex and by age, with the first principal component of the data largely separating the samples by sex (28% variance) and the second by age (23% variance) (**Figure 2a**). From the data, we first interrogated the contribution of sex factor to gene expression profiles that are consistent in young adult and early aging hearts. This analysis identified an almost equal number of genes (2,666; 1,651 higher in male; 1,015 higher in female) that are differential across sexes at 10% FDR as there were in the age comparisons (**Supplemental Data S1**). At a more conservative 5% FDR cutoff, 1,777 sex-dependent genes were found with 1,124 higher in male and 654 higher in female. An inspection of the top sex-associated genes show that they include genes residing on sex chromosomes including Xist, Eif2s3y, and Ddx3y; as well as participating in known sex-determination processes (**Figure 2b**). On a pathway level, we found that sex-differential genes appear to primarily cluster around metabolic pathways, including genes involved in pyruvate metabolism and TCA cycle, (P.adj: 9.4e–3), mitochondrial translation (P.adj: 2.5e–2), and gluconeogenesis (P.adj: 2.5e–2); as well as baseline differences in cardiac conduction, autophagy, and mitophagy pathways (**Figure 2c, Supplemental Figure S1a**).

**Figure 2.**
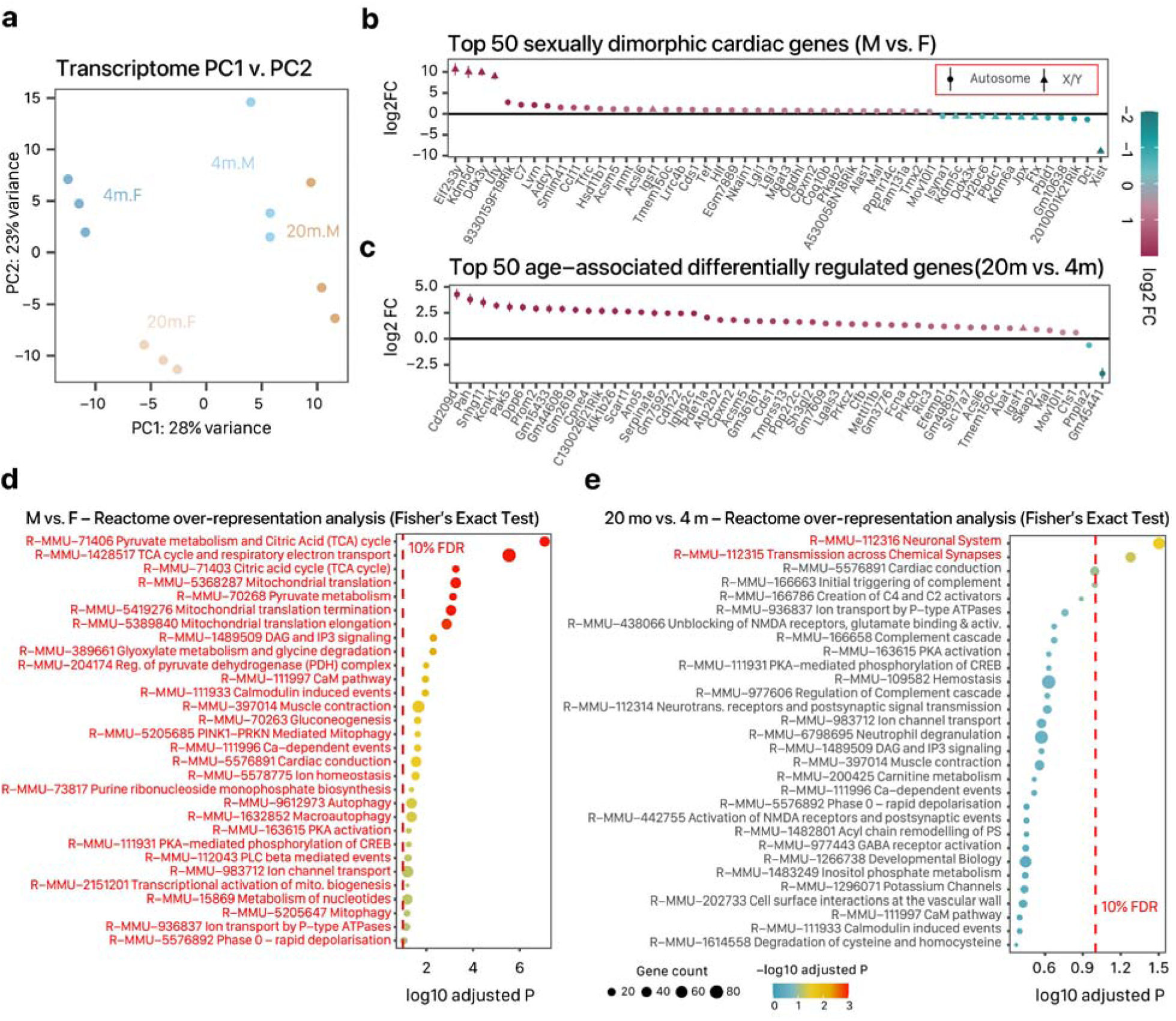
Age and sex associated cardiac genes. **a**. Scatter plot of first two principal components of total transcriptome profiles showing linear separation of samples by age and sex by grouping (n=3 per group). **b**. Point ranges of log2 fold changes and standard errors (y-axis) in male (M) vs. female (F) C57BL/6J hearts, showing the top 50 significant genes (x-axis). Point glyphs denote sex chromosome/autosome of the gene in mice. Color: log2 fold changes (FC). **c**. Point ranges of log2 fold changes and standard errors (y-axis) in sex-adjusted 20 mo. vs. 4 mo. C57BL/6J hearts, showing the top 50 significant genes (x-axis). **d**. Over-representation analysis against Reactome pathways (y-axis) among differentially expressed genes (10% FDR of |log2FC| ≥ 10%) in male (M) vs. female (F) hearts. Red dashed line: 10% FDR; point color: –log10 adjusted P; point size: number of quantified genes in a pathway. Red pathways are significant within 10% FDR. **e**. Over-representation against Reactome pathways (y-axis) among differentially expressed genes (10% FDR of |log2FC| ≥ 10%) in 20 mo. vs. 4 mo. hearts.

Upon adjusting for baseline sex differences, we further identified 2,519 genes that are significantly differentially expressed in 20 mo. vs. 4 mo. mouse hearts, with at least |log2FC| ≥ 10% at a conventional 10% FDR, including 1,464 up-regulated genes and 1,055 down-regulated genes. At a more conservative 5% FDR of |log2FC| ≥ 10% threshold, 1,738 genes were differentially expressed including 1,052 up-regulated and 686 down-regulated genes (**Supplemental Data S2**). We note that almost 40% fewer genes are found to be age-regulated when sex differences at the baseline are not taken into account (1611 vs. 2519 at 10% FDR), which would be consistent with significant sexual dimorphism in gene expression in both young and aged hearts and underlines the critical importance of taking sex into account in aging comparisons. The differentially expressed genes include 45 of 98 previously reported proteins found in aging hearts using microarrays (Bartling et al., 2019) including immune genes Cfb and Cd209d, whereas the most significantly changed genes also included several poorly characterized genes that are associated with age-associated diseases in other tissues (e.g., Ric3, Parkinson’s disease; Prkcz, cancer) (**Figure 2c; Supplemental Figure S1b**). Intriguingly, a conventional over-representation analysis using Fisher’s exact test against Reactome annotations found only two pathways to be enriched within the 1,738 (2,519) aging-differential genes (R-MMU-112316 Neuronal System and R-MMU-112315 Transmission Across Chemical Synapses) which have no obvious connection to cardiac aging processes (**Supplemental Figure S1a**). Similarly, we performed gene set enrichment analysis (GSEA) against MSigDB Canonical Pathways but likewise found few easily interpretable pathways (**Supplemental Figure S1b**), together suggesting although a sizeable number of age-differential genes are found in the model, they do not readily map to known processes through conventional enrichment comparisons (see below).

### Sex-dependent aging genes are enriched in calmodulin signaling

To discover gene signatures of sex-specific differences in aging, we next performed differential gene expression analysis on the age–sex interaction term and found 853 genes with sex-specific response to aging proceseses (336 higher in male:aged, 517 lower) at 10% FDR (553 genes with 234 higher in male:aged and 319 lower at 5% FDR) (**Figure 3a; Supplemental Data S3**). Among these genes are *Sprr1a* (small proline rich protein 1a), which is induced in aging more highly in male than female; *Cd72*, which encodes a protein functioning in B-cell proliferation and which is induced in aging in male but suppressed in female; *Tfrc*, which mediates iron update and is elevated in female in aging but reduced in male; and *Cpxm2*, which encodes a carboxypeptidase and which is induced more prominent in aging in female than in male (**Figure 3b**). A functional analysis using Fisher’s exact test for genes differentially regulated at 10% FDR against Reactome annotation showed significant enrichment of two clusters of terms, including DAG and IP3 signaling (R-MMU-1489509; P.adjust 0.0013), Calmodulin induced events (R-MMU-111933; P.adjust 0.0030), and PKA activation (R-MMU-163615; P.adjust 0.0073) which are mediated by *Adcy1, Adcy5, Adcy9*, Prkce, and other genes (**Figure 3c**); as well as an enrichment in the mTOR signaling term (R-MMU-165159; P.adjust 0.032) though this enrichment is mediated by the translation initiation factors *Eif4g1 and Eif4e*, the AMPK subunit Prkaa2, and other genes not part of hte mTORC complex (**Figure 3c**). A complementary gene set enrichment analysis which does not rely on arbitrary P value cutoffs further reveals an enrichment in cell cycle (R-MMU-69278, P.adjust 0.013), mitochondrial translation (R-MMU-5368287, P.adjust 0.041), and neddylation (R-MMU-8951664; P.adjust 0.067) related genes (**Figure 3d**). Thus the age:sex interaction analysis revealed two types of patterns: among prominently dimorphic age-associated genes that are significant at an individual gene levels, there is a *relatively* lower induction (or more severe decline) in aging females than aging males in genes related to calmodulin signaling; whereas when the entire transcriptome is concerned, in female aging hearts there are unevenly distributed trends of less *apparent induction* (or trending more toward decline) of genes associated with cell cycle and mitochondrial translation.

**Figure 3.**
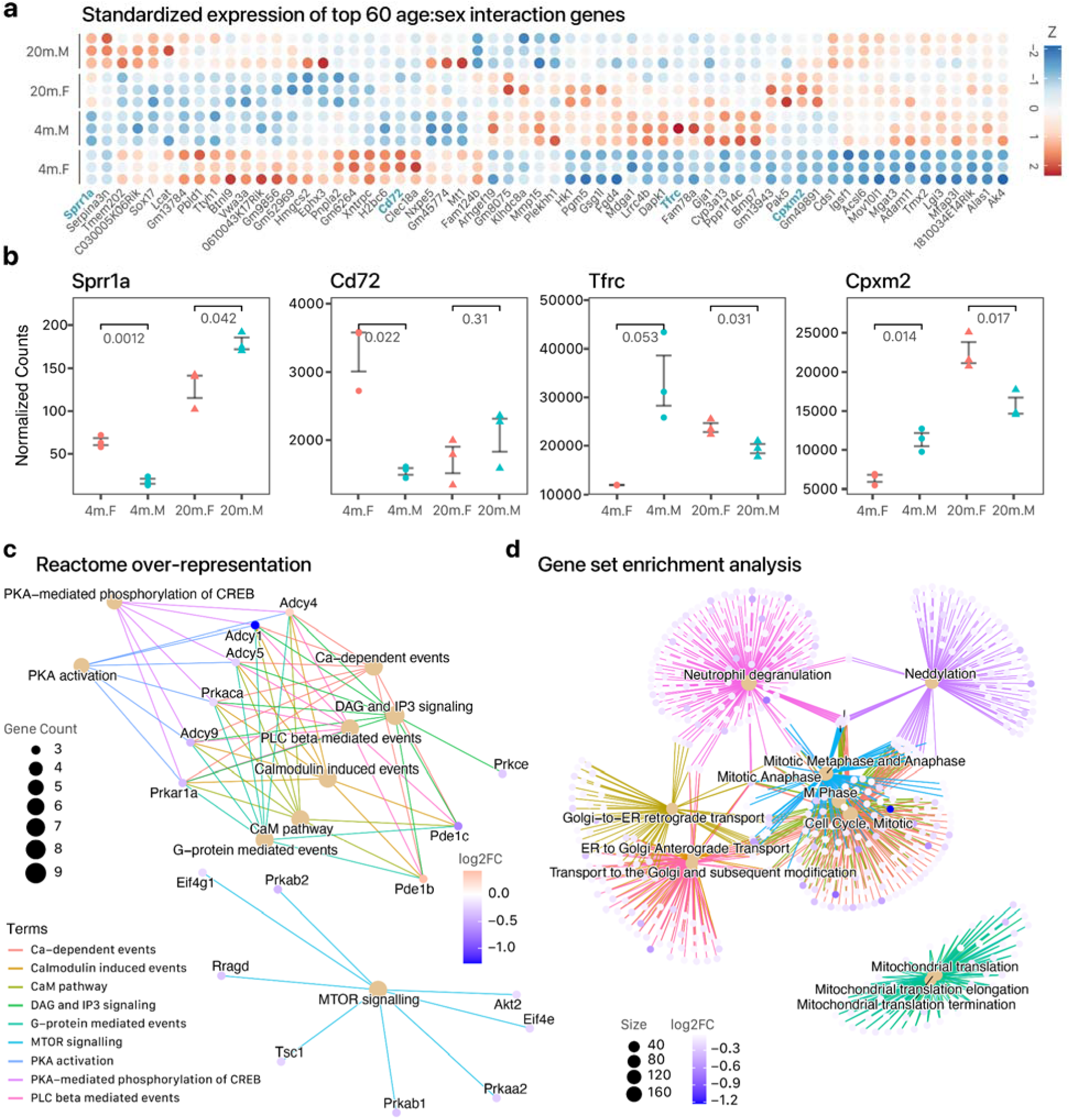
Cardiac genes and pathways with significant age:sex interaction. **a**. Heat maps showing the top 6 sexually dimorphic aging genes that show significant differences in aging response across sexes (DESeq2 age:sex 10% s-value). Colors denote standardized expression (Z score of normalized read counts). **b**. The normalized read counts of four genes from panel a were visualized across groups individually. P values: individual t-test in M vs. F comparison in 4 mo. and 20 mo. groups. **c**. Gene-concept network of two groups of significantly enriched Reactome terms among genes with significant age:sex interaction. Brown nodes are Reactome terms, linked through edges to their annotated genes in the foreground of the functional enrichment analysis. Gene node colors: log2 FC in male aged/adult vs. female aged/adult. Reactome term node size: gene count. Edge color: Reactome terms. **d**. Gene-concept network involving top enriched term in gene set enrichment analysis of the ordered gene list (by log2FC, male:aged vs. female:aged), linked to their annotated genes as in panel c.

### Proteogenomics comparison of cardiac gene expression by age and sex

We next analyzed matching cardiac samples with mass spectrometry to acquire proteomics data. The proteomics data are expected to diverge from RNA sequencing data in several important ways: on one hand, at present a typical mass spectrometry data set is able to access less depth than RNA sequencing and also carries distinct sources of technical variations than transcriptomics data. On the other hand, protein-level data provide unique information to reveal post-transcriptional regulation that likely lies closer to overt phenotypes. We collected both left and right ventricular samples for mass spectrometry and performed a general comparison of protein levels across age, sex, and chamber factor. Blood protein contamination is more severe at the protein level than the transcript level.

In total, we quantified 4,334 unique proteins identified at 1% FDR by UniProt accessions from 5 separate blocks of TMT experiments (**Supplemental Figure S2a–b**). In total, 764 proteins that show suggestive evidence of differential regulations in aging hearts after adjusting for sex and chamber (limma sex-adjusted P value ≤ 0.05; 733 cross-referenced to gene names), 249 of which was significant at 10% FDR (**Supplemental Figure S2c; Supplemental Data S4**). We likewise identified proteins that are sexually dimorphic including a number of proteins that are nominally found in the plasma but sampled in the cardiac samples, e.g., the male vs. female comparison found higher levels of murinoglobulin which is known to circulate at higher levels in male mouse plasma (Yamamoto et al., 1985) (**Supplemental Figure S2d; Supplemental Data S5**). In contrast to the mRNA data, protein-level information on cardiac aging most primarily revealed differential regulations along organellar configurations in the heart, including in mitochondria, extracellular space, and RBP-enriched nuclear speckles (**Figure 4a**). At an individual protein level, when contrasting 20 mo. and 4 mo. mouse hearts, 249 unique proteins consistently changed across age after adjusting for sex and chamber difference at 10% FDR; and 64 protein groups were consistently changed across sex. To understand the effect of aging on cardiac proteome, we further performed a subgroup analysis where 20 mo. male compared to 4 mo. male and 20 mo. females were compared to 4 mo. female without adjusting for age-sex interactions. We found 645 unique proteins to be differentially regulated at 10% in early aging male vs young adult male, but only 31 proteins in the female comparisons.

**Figure 4.**
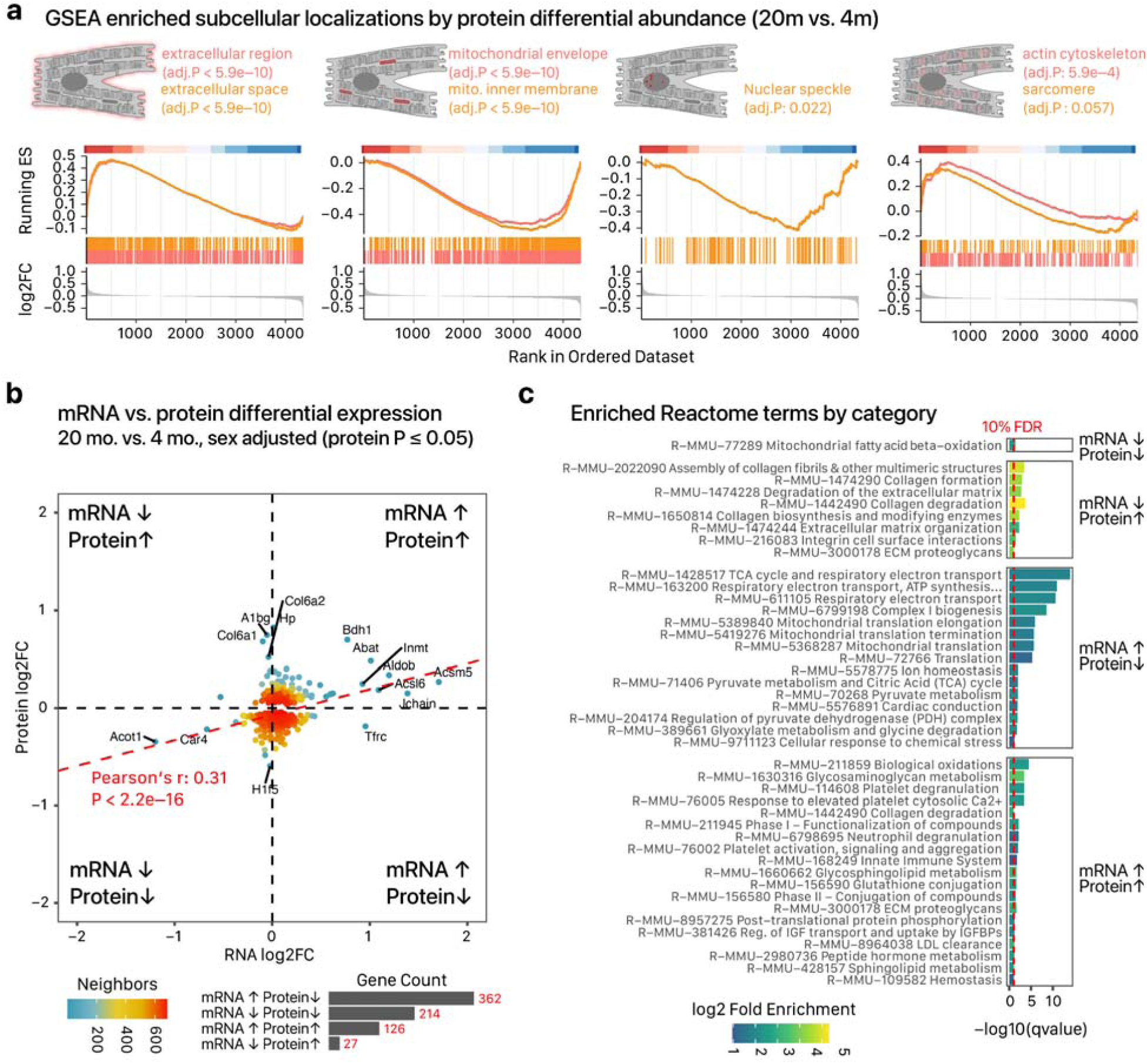
Comparisons of proteomics and transcriptomics data. **a**. Running enrichment score (ES) plot of significant subcellular localizations in gene set enrichment analysis (GSEA) of the quantitative mass spectrometry comparisons of 20 mo. vs. 4 mo. hearts. Proteins were ordered from highest to lowest log2FC. From left to right, age-associated proteins are enriched in extracellular (trending up-regulated), mitochondrial (down-regulated), nuclear speckle (down-regulated), and actin cytoskeletal (up-regulated) localizations. **b**. Scatter plot of log2FC among differentially regulated proteins (limma P ≤ 0.05) and the corresponding log2FC of their corresponding mRNA. Data are divided into four quadrants based on whether they show concordant or contradirectional protein and mRNA changes as labeled. Point colors: data density; bars: mRNA/protein counts in each quadrant. **c**. Bar charts showing significantly enriched Reactome terms (at 10% FDR) among proteins in each quadrant; x-axis: – log10 FDR-adjusted P value, Fisher’s exact test. Bar color: log2 fold enrichment against all quantified proteins.

Several observations were apparent in contrasting the mRNA-level and protein-level changes. First, there was minimal overlap between candidate differential proteins (limma P value ≤ 0.05) with significantly changed mRNA in the DESeq results (Jaccard index 0.08). Nevertheless, among these proteins we found a moderate but significant correlation between RNA and protein fold change (Pearson’s r = 0.31, P < 2.2e–16; Spearman’s rho 0.17, P 5.4e–6) consistent with previous reports (**Figure 4b**). In the sex-adjusted 20 mo. vs. 4 mo. comparison, as well as both subgroup analyses, we found myotilin, a protein involved in muscular dystrophy, to be significantly elevated with age (logFC 0.41 in both sexes, adj.P 0.022; logFC 0.53 in female, logFC 0.30 in male). Concordantly, myotilin is slightly up-regulated in the RNA level data (logFC 0.14 in both sexes) and is also upregulated from a previously published gene microarray data set that found 98 differentially regulated genes in 24 mo. vs. 6 mo. C57BL/6N mouse hearts (Bartling et al., 2019).

As expected, we observed more compressed fold changes at the protein level which are likely due to a combination of both technical reasons as well as the buffering effect due to the higher concentration of proteins than mRNA in a cell. A scatterplot of mRNA vs. protein fold change show that the commonly quantified mRNAs and proteins primarily occupied three quadrants based on their respective fold changes (**Figure 4b**); whereas down-regulation of RNA levels largely led to decreased protein levels, induction of mRNA can lead to either a corresponding increase of proteins or a paradoxical decrease of protein levels. Notably, the largest cluster involved contradirectional changes of increased mRNA but decreased proteins. Functional enrichment of the quadrant categories shows the non-diagonal quadrant (i.e., increased RNA but decreased proteins) to be enriched in large multiprotein complexes. Multi-protein complexes are known to show lower correlation between their mRNA and protein levels in part because of complex stoichiometry, i.e., an increase in one gene at the mRNA level would create an excess of a supernumerary subunit that does not necessarily lead to higher total number of functional complexes, thus creating a buffering effect that reduces the transmission of transcript-level regulation (**Figure 4c**). The results are consistent with an uncoupling of genetic regulations (induction of transcription) with the biochemical state (proteins) in the aging hearts. Moreover, they suggest that additional confirmations may be warranted for observed increases in age-associated mRNA encoding large complexes to corroborate their concordant changes at the protein level and hence potential downstream functional significance.

### Latent space representations reveal the involvement of RNA processing in aging

As the overrepresentation and gene set enrichment analyses unexpectedly returned few interpretable results in the sex-adjusted age comparison, we performed additional analyses to discern potential regulatory principles that are altered in the aging heart. To achieve this goal, we employed two complementary strategies to directly assess the concerted changes of extracted gene sets and modules. First, we performed gene set variance analysis (GSVA) (Hänzelmann et al., 2013), an unsupervised method that first calculates a gene set enrichment score for each sample and from there performs direct statistic testing of cross-sample score differences in multi-factorial experimental designs (i.e., ∽age + sex + age:sex). Using this approach, we found that mRNA processing and splicing pathways are heavily implicated in cardiac aging, with a total of ten Gene Ontology (GO) Biological Process (BP) terms enriched at 10% FDR, including BP: mRNA cis-Splicing via Spliceosome (P: 3.9e–4) and BP: mRNA Splice Site Selection (P: 4.9e–3) (**Figure 5a**). Other significantly enriched terms include BP: mRNA modification, BP: mRNA splice site selection, altogether suggesting the preferential involvement of module of mRNA processing and splicing related genes in sex-adjusted aging that evaded overrepresentation analysis. To corroborate the GSVA results, we further used a transfer learning strategy, where we first trained a model to learn the latent structure of mammalian gene expression using RNA sequencing data from over 17,000 human samples in the GTEx data set. We employed the matrix decomposition algorithm pathway level information extractor (PLIER) (Mao et al., 2019) to embed the gene-by-sample data matrix in a latent variable space, after which the algorithm learned the latent variables using the input gene expression data as well as a gene-pathway membership matrix. The pathway loading matrix learned from the large data set was then mapped to the current data set by assuming conservation of latent gene expression structure (Taroni et al., 2019) in order to assess the differential regulation of related gene modules in adult vs. early aging mouse hearts. Among the differential latent variables, we found two (Latent Variables 54 and 437) that were differentially abundant in aging at 10% FDR (P 1.1e–3 and 1.9e–4, respectively), and another related variable (Latent Variable 372) that was suggestively changed in the aging heart (P: 1.8e–2), to be related to pre-mRNA splicing and mRNA processing (**Figure 5b**). The top gene loadings associated with these latent variables confirm they map to various annotated mRNA splicing related terms including Reactome mRNA Processing, mRNA Splicing, Metabolism of RNA, and Spliceosome (Latent Variable 54); Reactome mRNA Processing (Latent Variable 372); and Processing of Capped Intron Containing Pre mRNA (Latent Variable 437) (**Figure 5c**). On an individual gene level, a number of transcripts encoding spliceosome and RNA-binding proteins were altered in aged hearts at 10% FDR including *Srrm1, Sf1, Srsf1, Srsf2, and Srsf7* (**Figure 5d**). Other known RNA-binding proteins that are altered in aging hearts include Rbm10 (logFC –0.18, s-value 9.9e–3), Rbm28 (logFC –0.18, s-value 0.040), and Ptbp2 (logFC – 0.19, s-value: 0.023) Taken together, these results from gene expression and functional analysis indicate that chronological aging exerts broad effects on mRNA splicing and processing pathways in the mouse heart.

**Figure 5:**
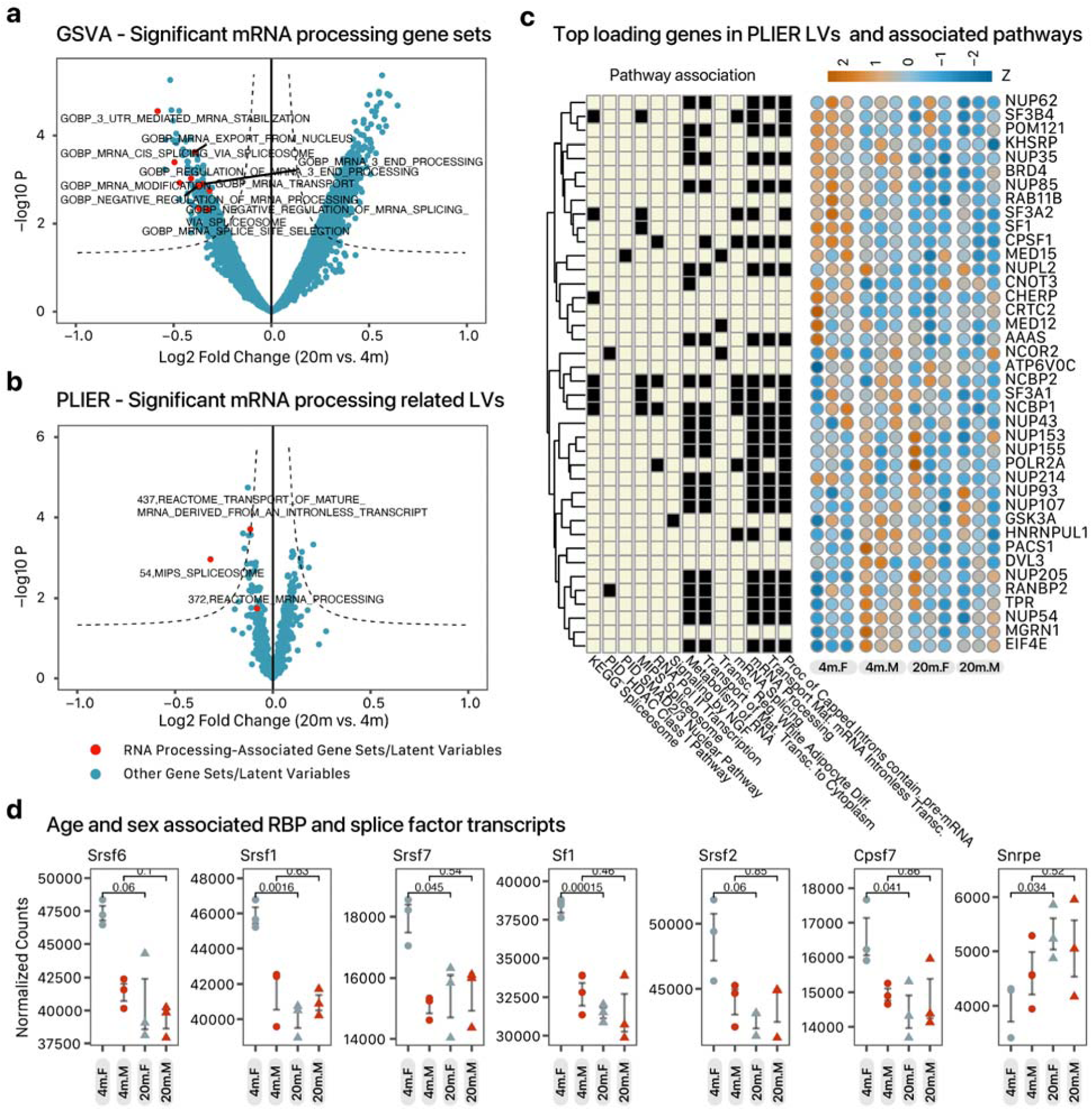
mRNA-processing and splicing related pathways in sex-adjusted aging hearts. **a**. Volcano plot showing the limma –log10 P value (y-axis) vs. log2 fold change (x-axis) of gene set variance analysis (GSVA) extracted gene sets. Gene sets related to mRNA splicing are labeled and colored in red. **b**. Volcano plot showing the limma –log10 P value (y-axis) vs. log2 fold change (x-axis) of PLIER-learned latent variables. Latent variables related to mRNA splicing are labeled and colored in red. **c**. Heat maps showing the top genes with highest loadings in the three mRNA splicing associated latent variables that are differentially regulated in aged hearts. Left matrix denotes membership of the gene in annotated pathways; colors in the right heat maps denote relative read counts in 20 mo. vs. 4 mo. female (F) and male (M) hearts. **d**. Individual plots of normalized gene counts across each group among selected significant transcripts encoding RNA-binding protein. P values: individual t-test between 20 mo. and 4 mo. hearts across age groups.

### Widespread reorganization of exon usage patterns in aging hearts

As the functional analysis implicates mRNA processing to be involved in aging, we next asked whether the changes in splicing related genes are associated with changes in alternative splicing patterns in the mouse heart. First, we used the statistical model in DEXSeq (Anders et al., 2012) to assess for the effects of individual exons within a gene and their interaction with age and sex factors on individual exon read counts. Differential exon usage can therefore be distinguished from differential gene expression by considering differential read counts between young adult and early aging hearts only in specific exons, which are first preprocessed into disjoint (non-overlapping) exonic parts for calculations in the DEXSeq model.

For example, a significant preferential decrease in the read counts of exonic parts 31–33 (adjusted P values: 1.2e–18, 8.9e–10, and 3.4e–4) in 20 mo. hearts vs. 4 mo. hearts can be seen in F-box only protein 11 (Fbxo11), a member of the F-box protein family whose members form parts of the RING-domain containing E3 ubiquitin ligase complexes (**Figure 6a**). The DEXSeq exonic parts 31–33 map through their genomic coordinates to the exon 1 of the Ensembl full-length Fbxo11-201 canonical transcript. Inference of the differential exon usage regions suggest the aging differences correspond to the Fbxo11-201 and Fbxo11-203 annotated protein coding transcripts, which are translated into UniProt accession Q7PD1-1 and Q7PD1-3, respectively, with the latter isoform omitting the majority of exon 1 and containing an alternative translation start site. The exon usage data is therefore consistent with a decrease in the shorter isoform of the FBXO11 protein in aging hearts. Another example is found in the polyhomeotic homolog 2 (Phc2), which is part of the large chromatin associated polycomb group multiprotein PRC1-like complex. Phc2 shows significant decrease in exon usage in exonic parts 19–21 and 28–29 (**Figure 6b**). The DEXSeq exonic parts 19–21 map to intron 9–10 in the Ensembl full-length Phc2-201 canonical transcript, and to the untranslated exon 1 of the shorter Phc2-202 isoform. Exonic parts 28–29 map to exon 10 of the full-length Phc2-201 isoform on Ensembl, and exon 2 of the shorter Phc2-202 isoform. Subsequent exonic parts mapping to other annotated exons (e.g., the DEXSeq disjoint exonic parts 32–35 mapping to the full-length Phc2-201 exon 11 and 12) were unchanged or slightly upregulated in aging hearts. Therefore although the RNA sequencing data strongly points to changes in exon usage patterns in the Phc2 gene, it is not possible from this analysis of collapsed short-read data to resolve whether the changes in exonic parts 19–21 and 28–29 corresponded to combinatorial differences in multiple annotated isoforms, or an unannotated age-specific events such as involving a cassette exon 10 or a partially retained intron. Nevertheless, the DEXSeq results strongly indicate widespread effects of exon membership on read counts suggesting a broad reorganization of transcript isoform patterns.

**Figure 6:**
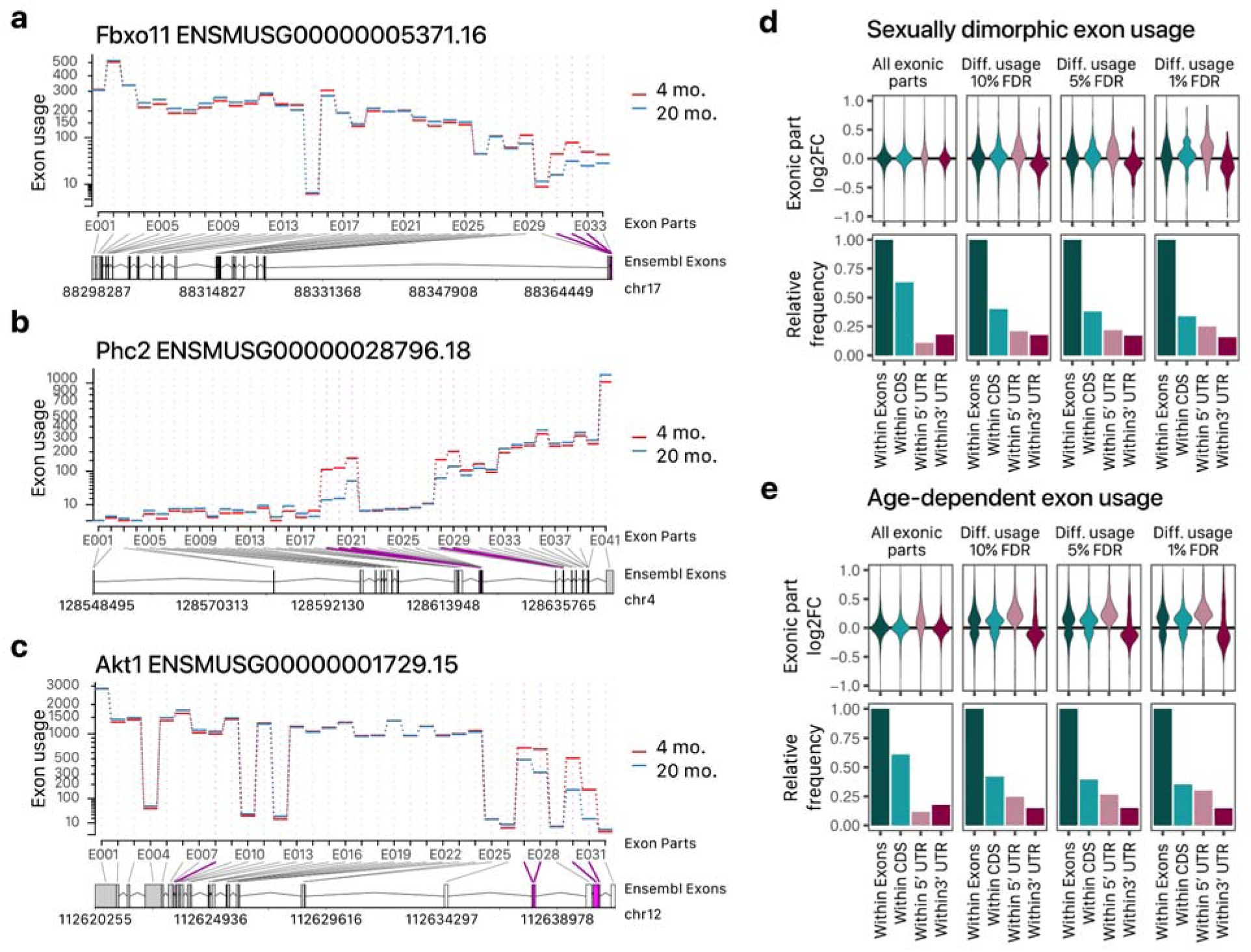
Differential exon usages in aging hearts. **a-c**. Exon usages in sex-adjusted comparisons between 20 mo. (blue) and 4 mo. (red) mouse hearts. Y-axis shows normalized read counts across disjoint exonic parts in DEXSeq after adjusting for gene-level differential expression. The disjoint exonic parts in DEXSeq are mapped to annotated exons in the Ensembl gene model below with the genomic coordinates labeled. Exonic parts with significant differential usage in 20 mo. vs. 4 mo. mouse hearts are colored in purple. **d**. Top: the distributions of exonic parts log2FC in male vs. female mouse hearts that overlap with different annotated gene regions in GENCODE annotations within each panel, from left to right: all exons, all CDS, 5’ UTR, and 3’ UTR. Subpanels denote all exonic parts, or those that show significant differential usages in male vs. female at different FDR cutoffs (from left to right, 10% FDR, 5% FDR, 1% FDR). Bottom: the relative frequency of exonic parts normalized to the total number of exonic parts within all GENCODE annotated exons. **e**. Distribution of lgo2FC and frequenc of exonic parts in sex-adjusted 20 mo. vs. 4 mo. mouse hearts. Subpanels are ordered as in panel c.

Sarcomeric genes are known to be under heavy influence of alternative splicing regulation, and we find complex and pervasive differential exon usage, including in **Myh7, Obscn, Myom1, Ttn**, and others (**Supplemental Figure S2a–S2b, Supplemental Table S2**). Other cardiac genes with known functional significance showed differential exon usages in the aging heart include Akt1 with lower coverage in aging in exonic parts 27, 28, 30, and 31 corresponding to a 5’ untranslated region of the full-length Akt1-201 canonical transcript in Ensembl (**Figure 6c**); as well as Sirt1 with lower coverage of DEXSeq exon part 16 in aging corresponding to the first translated exon (exon 6) of the canonical Sirt1-203 transcripts on Ensembl. In total, we found that age is associated with significant differences in exon usage of 7,520 exonic parts corresponding to 3,617 genes at 10% FDR. At a more conservative 5% FDR, there are significant exon usage differences in 4,238 exonic parts in the cardiac transcriptome belonging to 2,315 genes (**Supplemental Data S6**). Exonic parts with differential usages appeared to be unevenly distributed among coding regions with an enrichment for 5’ FDR overlapping exons (**Figure 6d–6e**).

Notably, integrated analysis of mRNA abundance and exon usage reveals that a greater number of genes are consistently affected by aging by way of exon usage pattern and splicing than those that are differentially expressed across identical significance thresholds. Moreover, genes that exhibit differential exon usage across age appear to overlap only minimally (214 or 9%) than differentially expressed genes, suggesting differential exon usage is in a position to exert independent effects from differential gene-level expression to aging hearts. Overrepresentation and network analyses suggest that genes with differential exon usage in aging hearts are clustered along specific functional categories including RNA metabolism, ribosome, nonsense mediated decay (NMD), as well as mitochondrial metabolism pathways (**Supplemental Figure S4a-b**). The overrepresentation of RNA metabolism terms suggests a possibility of positive feedback in transcript processing in aging where initial changes in RNA-binding proteins and splicing factors cause further changes in transcripts involved in post-transcriptional regulation processes. Taken together, these results are consistent with previous observations that transcript splicing may be more prominently altered in aging than gene-level expression in GTEx data (Wang et al., 2018) although they contrast somewhat with recent findings in human skeletal muscle aging where relatively few (163 total) transcripts showed differential exon usage at 10% FDR.

We likewise observed differences in exon usage across sex, affecting 1,537 exon parts in 1,056 genes by sex, and 1,868 exon parts in 1,168 genes by age-sex interactions (S**upplemental Data S7–S8**). Genes with sex differential exon usages included the 5’ exons in *Obscn and Pdlim5* (**Supplemental Figure S4c**). The first two 5’ exons in Mt1 display a complex age-sex interaction, with exon usage increasing during female aging hearts but decreasing in male aging hearts. Again we observed a poor overlap between differential exon usage genes and differential expression genes in se and age-sex interaction comparisons (171 genes or 16% in sex comparison, 48 genes or 4% in age-sex interaction), further highlighting that exon usage constitute an independent feature of sex-specific gene regulation. Taken together, the results from differential expression and exon usage analyses corroborate that changes in RNA processing and alternative splicing present a salient feature in cardiac aging processes.

### Detectability of alternative transcripts at the protein level

Finally, we asked whether any differential exon usage might show discernible proteome-level consequences, which would be important in establishing their molecular and potential functional significance. The extent to which alternative transcripts influence the composition of proteomes remains poorly understood. Alternative splicing can influence proteome compositions in at least two ways, first by diverting total transcript usage toward or away from productive, canonical isoforms to modulate total protein levels, and second by the production of transcript isoforms encoding different protein primary sequences (splice proteoforms). It is now well established that many alternative isoforms contain premature termination codons and so are likely subjected to nonsense-mediated decay, whereas others may produce nascent peptides that are degraded co-translationally, hence the extent to which alternative splicing events may influence cardiac functions await further elucidation at the protein level.

Alternative protein isoform identification in shotgun proteomics remains highly challenging for a number of reason, including the low abundance and poor annotations of non-canonical proteins, the requirement to detect a specific isoform-typic peptide not shared from the canonical sequence, and the prevalence of tryptic lysine residue around splice junctions. To examine whether the identified splice events may translate to stable proteins at the proteome level, we used a proteogenomics approach based on RNA-seq derived protein sequence database to re-analyze the acquired mouse heart proteomics data. Because common mouse proteomics databases remain incompletely annotated with regard to alternative splice isoforms (e.g., UniProt SwissProt contains 8,318 non-canonical sequences), this workflow allows potential undocumented isoforms to be salvagable from mass spectrometry data while avoiding runaway database size inflation and the accompanying false positive identifications.

To do so, we used a computational pipeline we previously developed to filter detected splice junctions in the RNA sequencing data and write likely detectable alternative protein sequences in custom protein sequence FASTA files. We first modeled the potential alternative splicing events that can explain the exon usage in the RNA sequencing data through exon splice junction spanning reads using rMATS v.4.1.0. This allowed splice junction spanning reads to be identified to model splice events including skipped exons (SE), mutually exclusive exons (MXE), alternative splice donor/acceptor sites (A5SS/A3SS), and retained intron (RI) events. A comparison of exon inclusion levels from these splicing events performed on two-sample aged vs. young adult male hearts and aged vs. young adult female hearts separately confirmed that splice event changes are prevalent, with 2862 splice events in both comparisons belonging to a total of 2911 genes to show significant changes in aging hearts at 10% FDR, among which 977 overlapped with genes with significant differential exon usages.

Secondly, this analysis also confirmed that a majority of alternative splicing events are likely undetectable as protein, both due to the very low transcript count and because they introduce frameshift leading to premature termination, although a substantial number of potentially translatable splice isoforms remain (**Figure 7a**). In the former case, we note that the majority of modeled splicing events in the cardiac transcriptome can be categorized as cassette exons (skipped exons). A large number of cassette exons have very low read counts and are thought to arise from splicing noise (Wan and Larson, 2018) and thus we reason they are unlikely to be translated into functional protein products or even so exist at a level detectable by mass spectrometry-based proteomics. The splice junction read count distribution can be modeled using a gamma-gaussian mixture model (**Figure 7a**) in RNA-seq data to separate the splice junctions into low and high read populations, which allows the opportunity to compare translation proportion or alternatively to remove primarily low-count skipped exons that fall into a distinct distribution of read counts from the majority of other exon junctions when generating RNA-guided protein sequence databases (see Methods).

**Figure 7:**
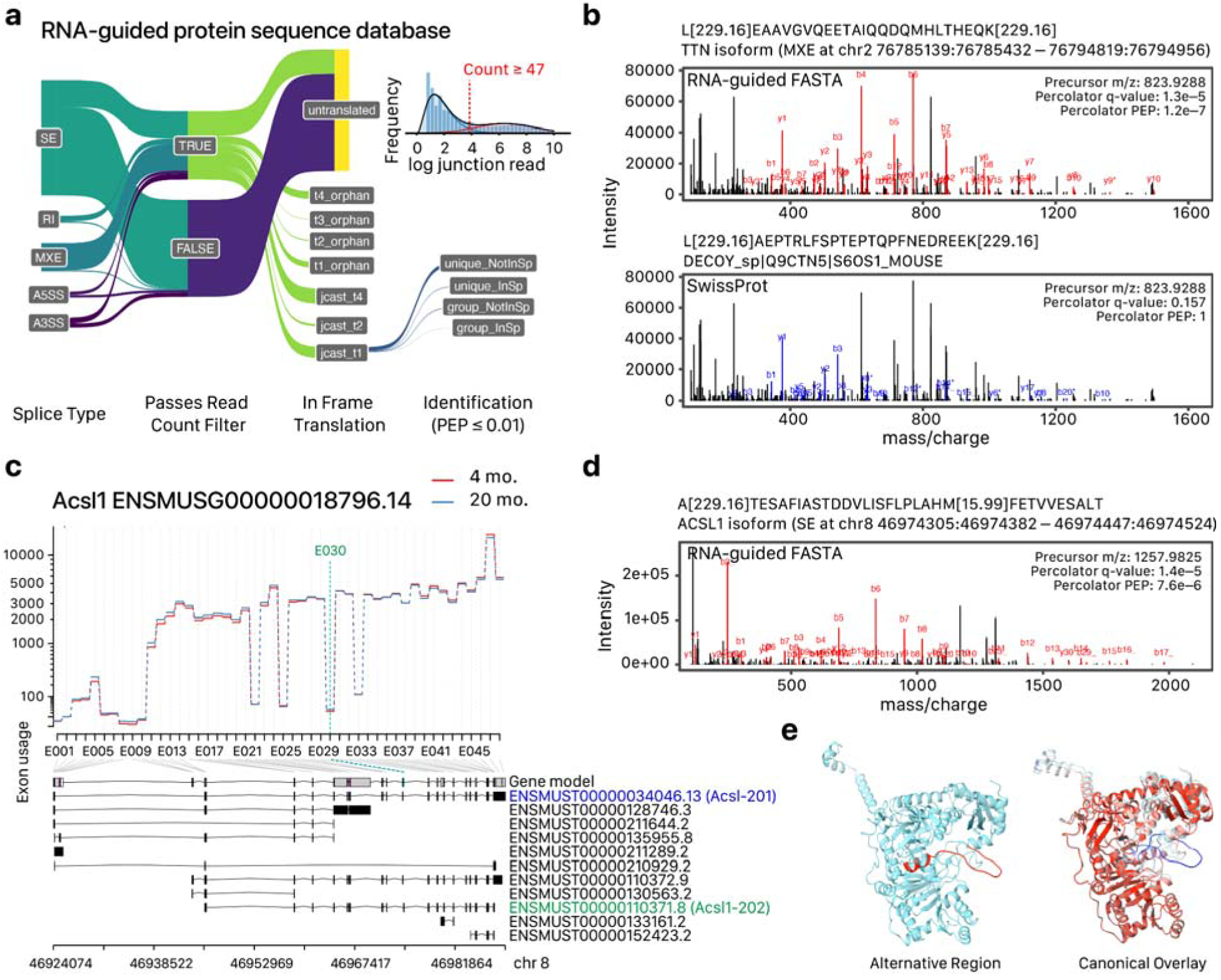
Protein-level consequences of exon usage changes. **a**. Sankey diagram illustrating the fate of all modeled splice junctions in the RNA-guided protein database methods. Node sets from left to right denote the modeled splice type of the exon junctions, whether the skipped junction count is above a threshold modeled from total junction read counts, translation status (see Methods), and identification status. Inset: modeled read count cutoff. **b**. Top: peptide spectrum match (PSM) of a non-canonical peptide belonging to titin from a database search using the RNA-guided protein database in panel a. Matched peptide fragment b- and y-ions are labeled and colored in red. PEP: Percolator PSM posterior error probability. Bottom: the best-hit candidate sequence matched to the identical spectrum as the top panel when queried against UniProt SwissProt canonical and isoform entries. **c**. Differential exon usage of Acsl1 in 20 mo. vs. 4 mo. mouse heart. An disjoint exonic part E030 (green) shows higher usage in aged hearts and maps to an exon absent in the canonical transcript (Acsl1-201). **d**. Peptide-spectrum match of a non-canonical peptide corresponding to a cassette exon matching to E030 identified in the quantitative mass spectrometry data using the RNA-guided protein database. **e**. Predicted protein structure of the predicted full-length alternative ACSL1 containing the translated alternative region from which the non-canonical peptide was identified, joined to the full length UniProt canonical sequence via a 10-amino acid joint in the upstream and downstream exons. Left: predicted protein structure with the alternative region highlighted, right: the predicted structure of the alternative region overlaid on the predicted structure of the canonical protein.

From the custom databases, we identified 1,812 peptide sequences at 1% Percolator PEP that are uniquely mappable within the FASTA database to non-canonical protein isoforms, 1,446 of which are not found among UniProt SwissProt mouse isoform sequences at the time of writing. In addition, 253 potential canonical isoforms are potentially identifiable as part of a unique protein group that does not include a canonical sequence, 170 of which are not found among UniProt SwissProt mouse isoform sequences at the time of writing. The identification of non-canonical peptides demands caution as there are potential alternative explanations of peptide spectrum matches such as from unaccounted canonical peptides or modifications. We manually examined a subset of identification and found that at least a portion of non-canonical ID may be detectable. For instance, we identified a titin isoform peptide which is modeled to originate from a mutually exclusive exon and which is not mappable to UniProt SwissProt or Trembl sequences at the time of writing (**Figure 7b**), which suggests novel protein isoforms remain to be found in the cardiac proteome, although safeguard measures will be needed to exclude common sources of spurious identification (see examples below).

From the labeled quantitative proteomics data we further attempted to quantify the differential abundance of non-canonical sequences across age and sex groups. There are multiple challenges associated with quantifying isoform peptides, including the difficulty of identifying these sequences confidently and the high burden of multiple testing. Moreover, the current workflow does not incorporate match between runs, as the false discovery rates of FBRs remain unclear and can compound the caveats of non-canonical sequences. Accordingly, we were able to gather age:sex quantitative information on 115 translated non-canonical isoform groups in the heart belonging to 86 genes, which were identifiable at 1% posterior error probability in at least 1 of 5 blocks of TMT experiments and for which sufficient samples remained for 20mo. vs 4mo. comparisons in male and female hearts (**Figure 7c**). The identified alternative isoforms map to multiple cellular compartments, suggesting alternative splicing has the potential to exert far reaching consequences on cardiac proteome makeup and function. We then compared age associated expression in old vs. young males and old vs. young females. When comparing the TMT channel intensity data, we found 15 unique isoform peptides to show suggestive evidence of differential expression in age at limma P ≤ 0.05 in male and 11 in female, although only 8 in male and 2 in female reached significance following multiple testing corrections at 10% FDR. The potential differential isoform peptides include those belonging to known RBM target genes, including Ldb3 (An SE peptide, logFC 0.39, adjP 0.046 in male) and Mlip (logFC 0.34, adj P 0.051 in male). We examine here two potential cases where transcripts with differential exon usage may be linked to protein-level changes.

In the second example, we identified a putative isoform peptide from Acsl1. Acsl1 codes for the long-chain fatty-acid-CoA ligase 1 which participates in beta oxidation and is found in the mitochondria and the peroxisome. At the transcript level, exon usage analysis shows a complex pattern of age vs. young patterns across disjoint exon parts suggestive of complex splicing pattern. We focused on exon part 30 which shows a suggestive increase in exon usage in aging hearts after adjusting for sex (**Figure 7c**). Splice junction modeling predicts a skipped exon event where a cassette exon (chr8:46974447– 46974524) is optionally included in between the upstream (chr8:46974305–46974382) and downstream (chr8:46977980–46978114) exons. The exon triads correspond to Ensembl canonical transcript (Acsl-201) exons 11-x-12, with the cassette exons not found in the canonical transcript but part of an APPRIS alternative transcript (Acsl-202) in the gene model where the triad corresponds to exons x-10-11. From the proteomics data, we tentatively identified a peptide that corresponds to the in-frame translation product of a transcript linking the upstream Acsl-201 exon 11 with the cassette exon corresponding to alternative Acsl-202 exon 10, resulting in a peptide not found in TrEMBL. This peptide sequence is 33-residue long and semi-tryptic A[229.16]TESAFIASTDDVLISFLPLAHM[15.99]FETVVESALT with moderate spectral evidence (**Figure 7d**), hence a false discovery identification of the peptide cannot be ruled out, but we note that the peptide sequence was identified at least 12 times across 5 blocks of TMT quantitative proteomics data at percolator PEP ≤ 0.01 (median percolator PEP of 0.00028; max 7.6e–3; min 7.6e–6), with a number of diagnostic fragments (y1, y2, y3, y4, y6, y7, y9, y12, y20, y1_, y2_, y3_, y10_, y30_) that distinguish the variant portion of the sequence from its canonical-invariant portion in the Acsl1 gene (ATESAFIASTDDVLISFLPLAHMFETVVE[…]) (**Supplemental Figure S5a–S5b**). In the UniProt SwissProt canonical/isoform database search the top scoring spectrum was non-identifiable to a forward target sequence and was assigned to a decoy sequence (**Supplemental Figure S5c**), suggesting the isoform database afforded substantial increase in power to explain the spectrum by peptide spectrum match. To explore whether the spectrum may arise from unknown modifications, we further performed an open mass window search using MSFragger v.3.2 (Kong et al., 2017) followed by Percolator v.3.0. The spectrum was likewise unidentifiable and was assigned to n[229.1629]EATYGERVVAFAAVEGIFFSGSFASIFWLK[229.1629] sp|P11157|RIR2_MOUSE with a non-significant Percolator PEP (0.299), with none of the top 5 MSFragger PSM candidates matched to Acsl1. In the quantitative proteomics data, the putative protein isoform had a corresponding increase in aging (logFC 0.43, P Value 1.7e–3, P.adjust 0.078). Taken together we consider that there are multiple lines of tentative evidence that suggest the differential exon usage at a minor exon in Acsl1 has protein-level consequences. Intriguingly, the alternative peptide overlaps with the main AMP-binding motif in the ACSL1 protein. The alternative sequence is not predicted to be intrinsically disordered in either the deep learning based algorithm Metapredict v.2.0 and biophysics-based algorithms (IUPred3/ANCHOR2) (**Supplemental Figure S5d–e**). To corroborate the sequence analysis, we performed structural predictions on the modeled alternative full-length protein isoform using AlphaFold2, after the translated exon triads were joined back to the canonical SwissProt isoform sequence. The predicted protein folding suggests that the isoform-variant sequence belongs to a pocket of unstructured loop within the AMP binding domain that does not alter overall protein structure when overlaid on the canonical sequence structural model (**Figure 7e**).

In a second example, we considered the exon usage patterns of aspartyl β-hydroxylase (*Asph*). In mouse and in human, the *Asph* locus is known to contain multiple genes through alternative transcript processing or splicing: the 24-exon BAH, the 14-exon catalytically inactive humbug, and the 5-exon junctin (Dinchuk et al., 2000, 2002), as well as multiple isoforms of cardiac junctate (Treves et al., 2000), each with distinct molecular function and regulation. In the DEXSeq results, we saw multiple contiguous disjoint exonic parts (E028–E030) that were suppressed in aged hearts. Splice junction modeling revealed multiple MXE and SE events in the region that involved the optional inclusion for exonic part E030 corresponding to chr4:9624359–9624401 (**Supplemental Figure S6a**). This exonic part corresponds to the an exon between exon 3 and 4 of the APPRIS canonical Asph-202 transcript that codes for full-length ASPH/BAH, but does not constitute part of the canonical transcript but and a constitutive peptide of Asph-206/Trembl Q9CR06 (Junctin) and other transcripts in the *Asph* locus. From the mass spectrometry data, we identified two corresponding peptides LLEGPGGLAK, VLLEGPGGLAK (PSM Percolator PEP ≥7.4e–5) (**Supplemental Figure S6b–S6c**) which span exon parts E030 and E035. The peptides are part of SwissProt isoform but not in SwissProt canonical sequences (**Supplemental Figure S6d**). In the quantitative proteomics data, the isoform is significantly down-regulated in aged hearts consistent with the transcript data (protein logFC –0.22, limma P Value 4.5e–6, P.adjust 9.7e–4). Due to the nature of short-read sequencing and bottom-up proteomics data, it is not possible to discern which of the known full-length transcripts is differentially regulated. Nevertheless, the data are consistent with a rerouting of exon usage in aging hearts from the full length ASPH/BAH isoforms toward exons shared by *junctin/junctate* through splicing changes, which is likely transmitted to the protein-level.

## Discussion

Recent work has pointed to the intrinsic sexual dimorphism in cardiac aging processes in rodents and in humans as a modulating factor of cardiovascular risk. It has been well established that many cardiac genes have sex-differential expression patterns. Previous studies have found tens to hundreds of sexually dimorphic genes in the mouse heart (Isensee et al., 2008; Li et al., 2017; Tsuji et al., 2017), rat (Naqvi et al., 2019; Trexler et al., 2017), and in human hearts (Fermin et al., 2008; Gershoni and Pietrokovski, 2017; Isensee et al., 2008; Lopes-Ramos et al., 2020; Mayne et al., 2016; Naqvi et al., 2019; Newman et al., 2017) which involve both sex chromosome and autosomal genes participating in diverse processes particularly metabolism and ion transport pathways (Fermin et al., 2008; Naqvi et al., 2019); moreover, sex-biased gene expression may show sex-by-environment interactions with pathophysiological factors including cardiac hypertrophy, cardiomyopathies, and heart failure (Newman et al., 2017; Tsuji et al., 2017). In the present work, we examined cardiac gene expression across sexes and two age groups in C57BL6/J mice. The results show a large number of sex and age associated genes in the heart, and moreover that sex also exerts an influence on age-associated differential expression of multiple genes including those participating in camodulin signaling pathways.

Cardiac genes are prominently regulated through alternative splicing (Baralle and Giudice, 2017). Cardiac development and maturation involve successive changes in splice patterns modulated by the RBFOX, RBM, MNBL, and other families of splice factors, whereas the adult splicing program is reversed to the fetal/neonatal splicing pattern during pathological cardiac remodeling (Baralle and Giudice, 2017; Gao and Wang, 2020). In parallel, recent work has implicated changes in alternative splicing in the blood, the brain, and other tissues during organismal aging processes (Balliu et al., 2019; Deschênes and Chabot, 2017; Rodríguez et al., 2016; Stegeman and Weake, 2017; Wang et al., 2018). Despite progress, how alternative splicing in the heart varies by the interaction of age and sex remains incompletely understood and may shed light on established sex biases in the risks of age-associated heart diseases. Our analysis of sex-adjusted aging changes point to RNA-binding proteins and splicing factors as principally different in transcript levels in sex-adjusted aging hearts. These changes occur concomitantly with widespread exon usage differences in the cardiac genome, including in critical processes pertaining to post-transcriptional regulations, protein translation, and mitochondrial metabolism that are commonly implicated in heart diseases.

An outstanding challenge of alternative splicing research concerns examining the protein-level effects of exon usage, whereas current knowledge on which alternative transcript isoforms are translated into polypeptides and their molecular functions remains limited. Here we used an RNA-guided protein sequence database proteogenomics approach to identify splicing-derived proteoforms in mass spectrometry data. The results corroborate the present challenges of linking transcript level consequences of splicing to proteoform level changes. Further progress in this area will likely require continued advances in analytical and informatics approaches including more sensitive mass spectrometry, long-read sequencing (Miller et al., 2022) and top-down/middle-down proteomics methods (Melby et al., 2021). Nevertheless, with the present approach we were able to identify hundreds of non-canonical peptides that may be detectable from mass spectrometry data and that correspond to splice junctions modeled from the transcriptome profiles of young adult and early aging mouse hearts, including instances where exon usage changes at the the transcript level may be linkable to downstream protein-level consequences. Continued work to elucidate the interactions between RBPs, alternative splicing, and proteoforms orchestrate to alter cardiac proteome compositions and function will likely play an important role in advancing understanding of cardiac aging processes. Future work might also interrogate overlaps between the global remodeling of splicing patterns in cardiac again and those in pathological cardiac hypertrophy in order to identify potential correlates of age-associated risks of heart diseases.

## Supplemental Tables and Figures

**Supplemental Table S1:**
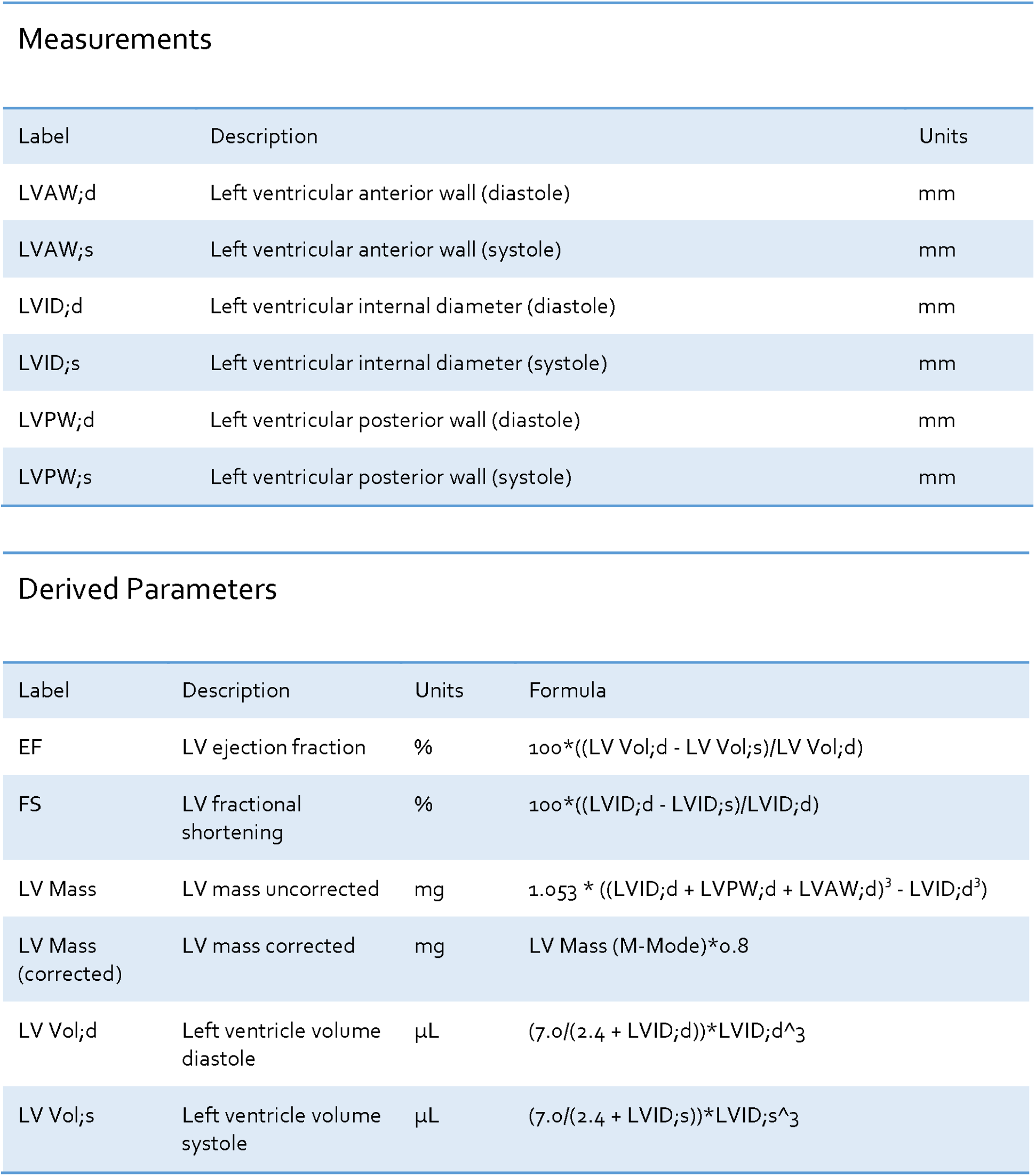
Measured and derived echocardiographic parameters.

Measurements
| Label | Description | Units |
| --- | --- | --- |
| LVAW;d | Left ventricular anterior wall (diastole) | mm |
| LVAW;s | Left ventricular anterior wall (systole) | mm |
| LVID;d | Left ventricular internal diameter (diastole) | mm |
| LVID;s | Left ventricular internal diameter (systole) | mm |
| LVPW;d | Left ventricular posterior wall (diastole) | mm |
| LVPW;s | Left ventricular posterior wall (systole) | mm |

| Label | Description | Units | Formula |
| --- | --- | --- | --- |
| EF | LV ejection fraction | % | $100 * ((LV\ Vol;d - LV\ Vol;s) / LV\ Vol;d)$ |
| FS | LV fractional shortening | % | $100 * ((LVID;d - LVID;s) / LVID;d)$ |
| LV Mass | LV mass uncorrected | mg | $1.053 * ((LVID;d + LVPW;d + LVAW;d)^3 - LVID;d^3)$ |
| LV Mass (corrected) | LV mass corrected | mg | $LV\ Mass\ (M-Mode) * 0.8$ |
| LV Vol;d | Left ventricle volume diastole | $\mu L$ | $(7.0 / (2.4 + LVID;d)) * LVID;d^3$ |
| LV Vol;s | Left ventricle volume systole | $\mu L$ | $(7.0 / (2.4 + LVID;s)) * LVID;s^3$ |

**Supplemental Figure S1:**
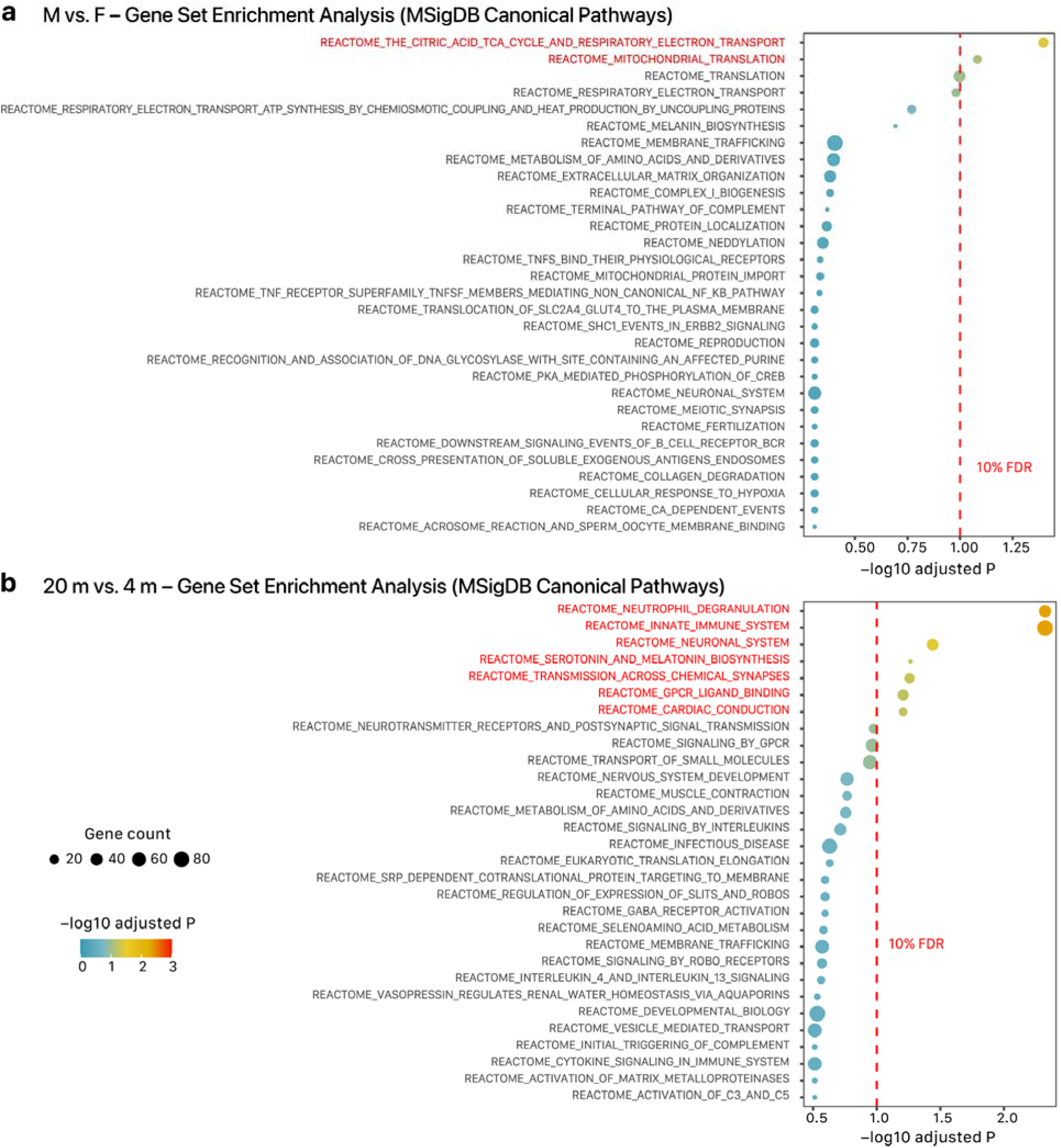
Gene set enrichment analyses of age and sex associated cardiac genes. **a**. GSEA of male vs. female mouse heart gene expression against MSigDB Canonical Pathways gene sets (fgsea permutation test with FDR-adjusted P ≤ 0.1; x-axis). **b**. Gene set enrichment analysis (GSEA) of 20 mo. vs. 4 mo. mouse heart gene expression against MSigDB Canonical Pathways gene sets likewise showed few interpretable gene sets (y-axis) with significant biased expression (fgsea permutation test with FDR-adjusted P ≤ 0.1; x-axis). Color denote –log10 adjusted P. Red gene sets are differentially regulated at 10% FDR.

**Supplemental Figure S2:**
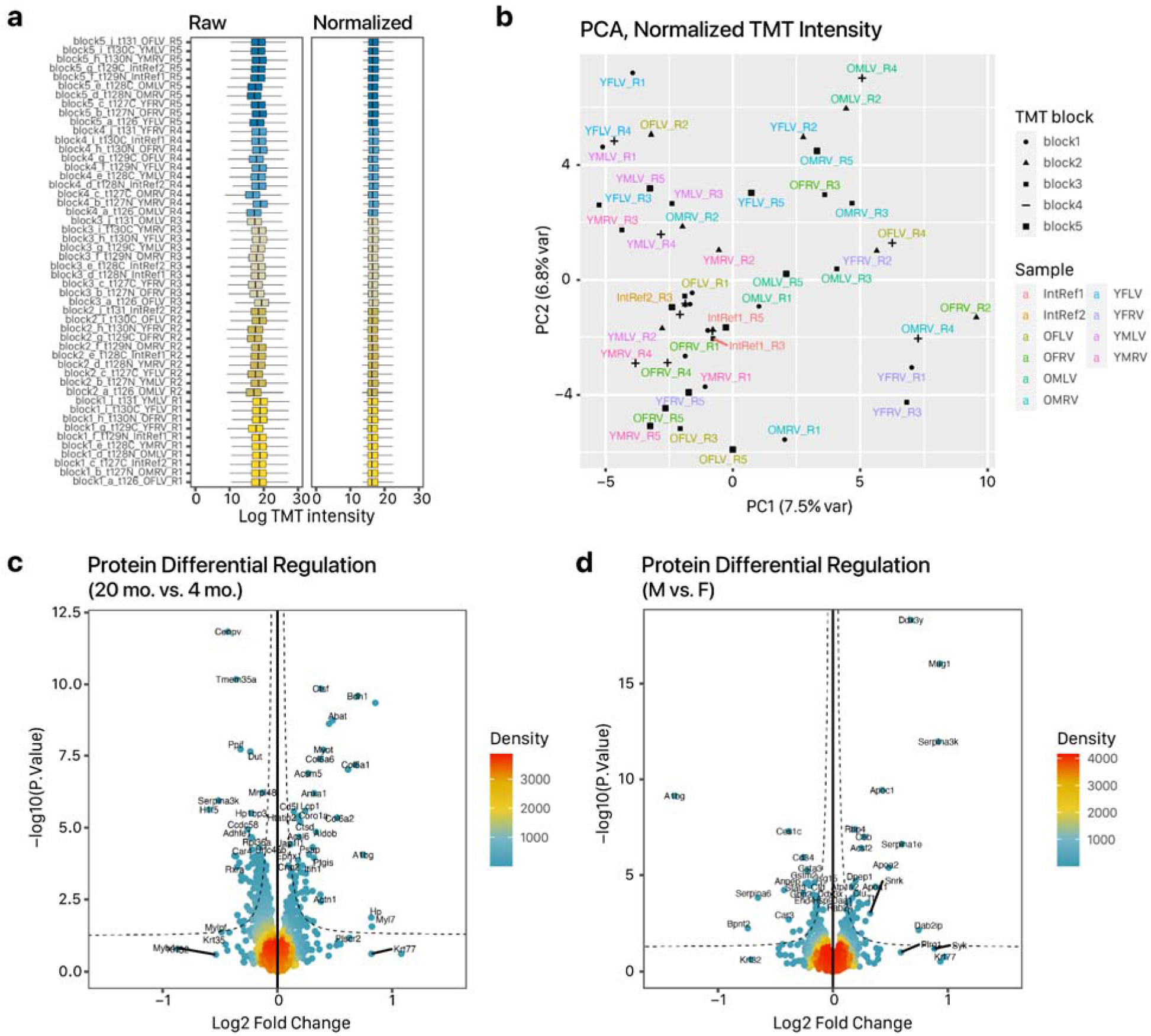
Proteomics characterization of age and sex differences in cardiac protein abundance. **a**. Box plots showing TMT label assignment across experimental blocks and TMT channels, and the corresponding samples. O: 20 mo. mouse heart; Y: 4 mo. mouse heart; LV: left ventricle; RV: right ventricle; M: male; F: female. Left: box plot of log raw TMT channel intensity. Right: log intensity following normalization. **b**. First two principal components of normalized TMT protein intensity values following normalization. Shape: experimental block; color: sample groups. **c**. Volcano plot of age comparison (20 mo. vs. 4 mo.) after adjusting for sex and chamber. X-axis: log2 fold change (20 mo. vs. 4 mo.); y-axis: –log10 limma P value. Color: point density. d. Volcano plot for sex comparison (M vs. F).

**Supplemental Figure S3:**
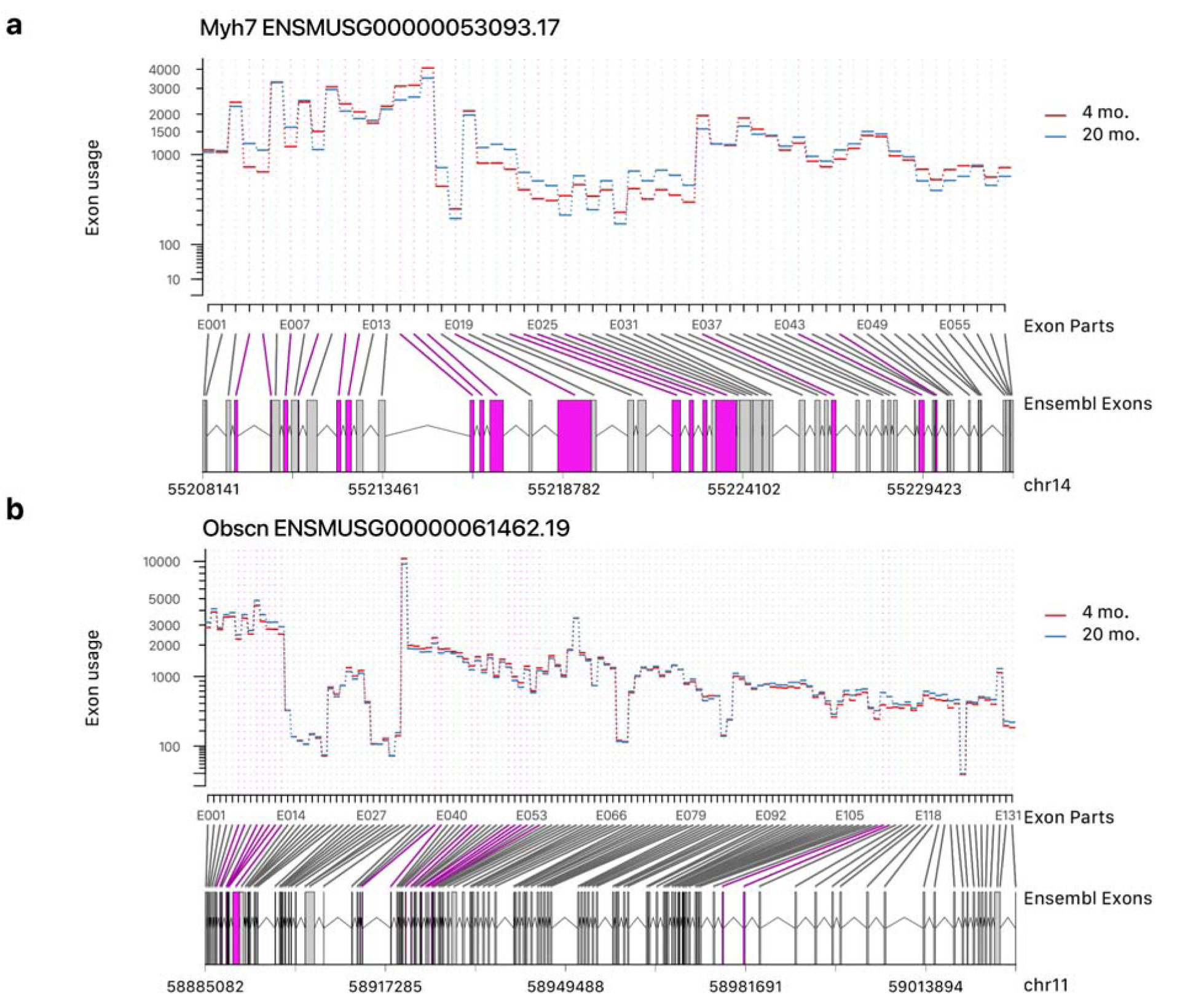
Age-associated exon usage patterns in sarcomeric genes. Exon usage graphs showing complex reorganization of age-associated exon usage patterns in two sarcomeric genes **a**. *Myh7* and **b**. *Obscn*. The DEXSeq disjoint exonic parts (x-axis) are mapped to Ensembl gene models and their genomic coordinates. Disjoint exonic parts with significant differences in exon usage are highlighted in purple.

**Supplemental Figure S4:**
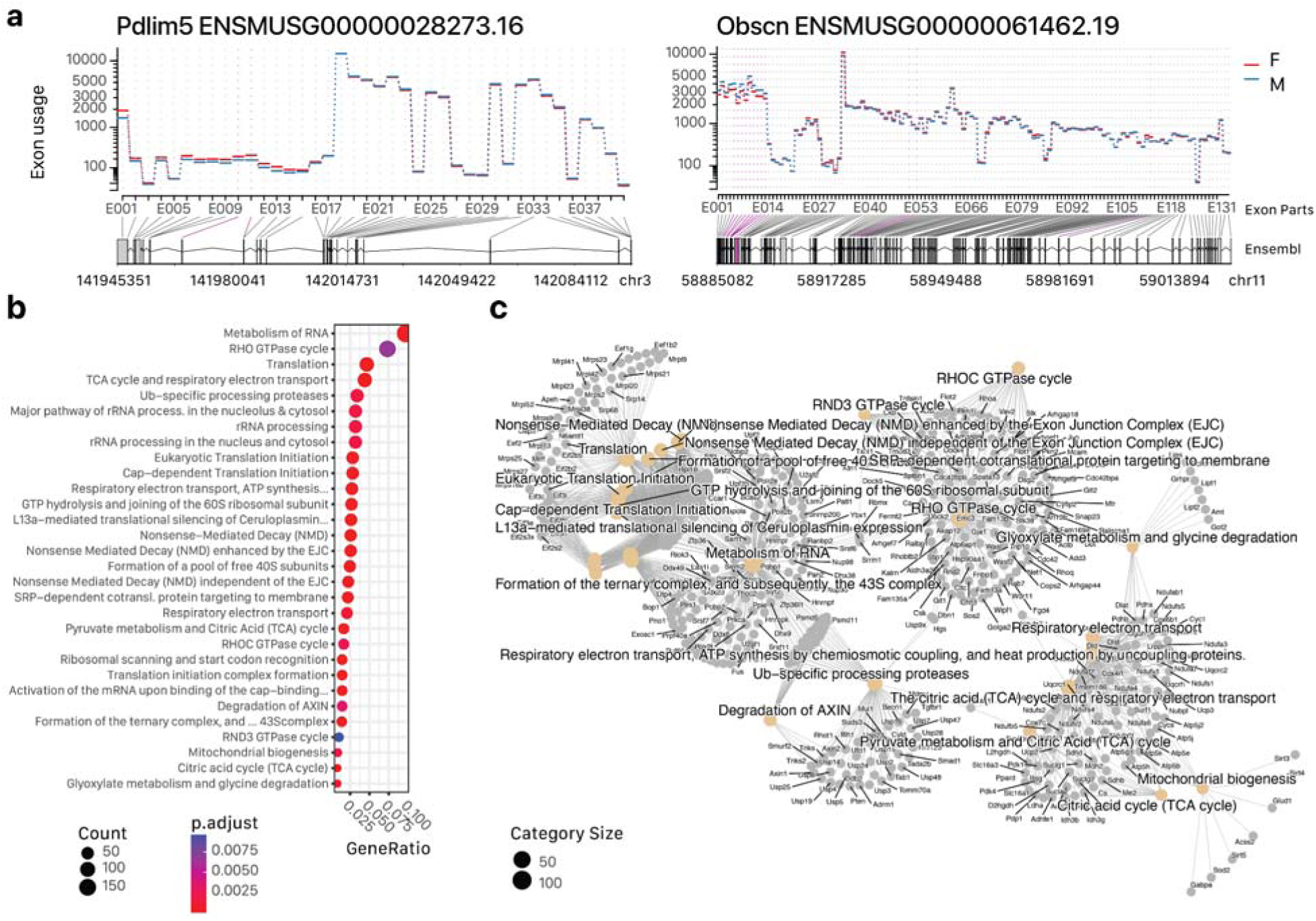
Age and sex associated exon usage differences in the heart. **a**. Examples of sex-dependent exon usage difference in two genes (Pdlim5 and Obscn). Disjoint exonic parts with significant differences across sex (M vs. F) are shown in purple. **b**. Dot plot showing the enrichment ratio and adjusted P values of over-represented Reactome pathways among genes with age-associated exon usage difference. Exon usage changes are enriched along RNA metabolism, ribosomal, and mitochondrial metabolic pathways. **c**. Gene concept network showing top 15 enriched Reactome terms and their associated genes with significant differences in exon usage in 20 mo. vs. 4 mo. hearts.

**Supplemental Figure S5:**
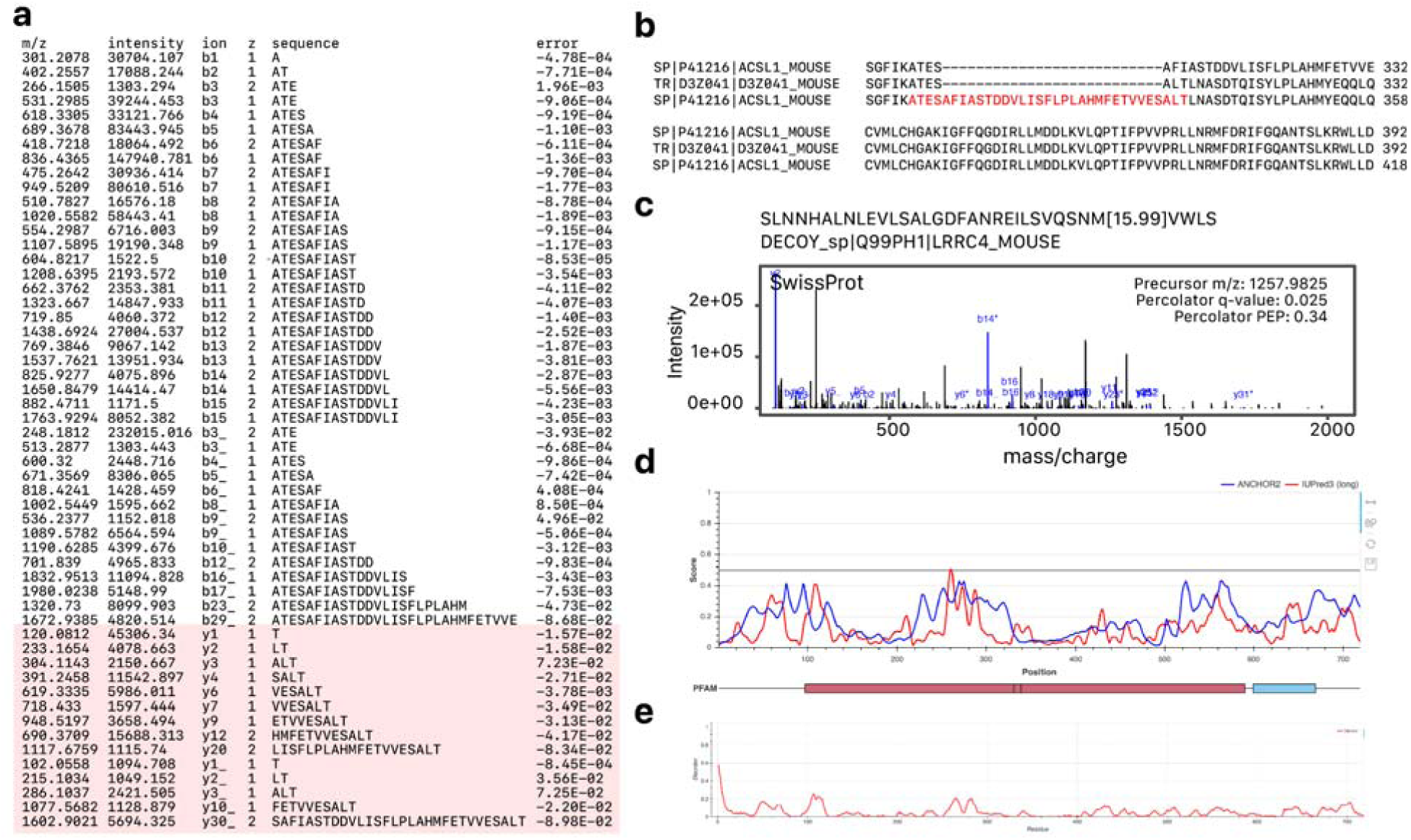
Putative identification of a non-canonical peptide in Acsl1. **a**. Matched fragment ions in the best-hit peptide spectrum match in the RNA-guided protein database query in the spectrum corresponding to Figure 7b. **b**. Sequence alignment in the region of interest in Acsl1 mapping to the alternative exon. The identified peptide sequence is highlighted in red. **c**. The best-hit peptide spectrum match in the UniProt SwissProt canonical + isoform database query in the same spectrum as Figure 7b. The spectrum was assigned to a decoy sequence SLNNHALNLEVLSALGDFANREILSVQSNM[15.99]VWLS belonging to DECOY_sp|Q99PH1|LRRC4_MOUSE, Percolator PEP 0.34) d. Predicted protein sequence disorder using ANCHOR2 and IUPRED3. **e**. Predicted protein sequence disorder using Metapredict.

**Supplemental Figure S6:**
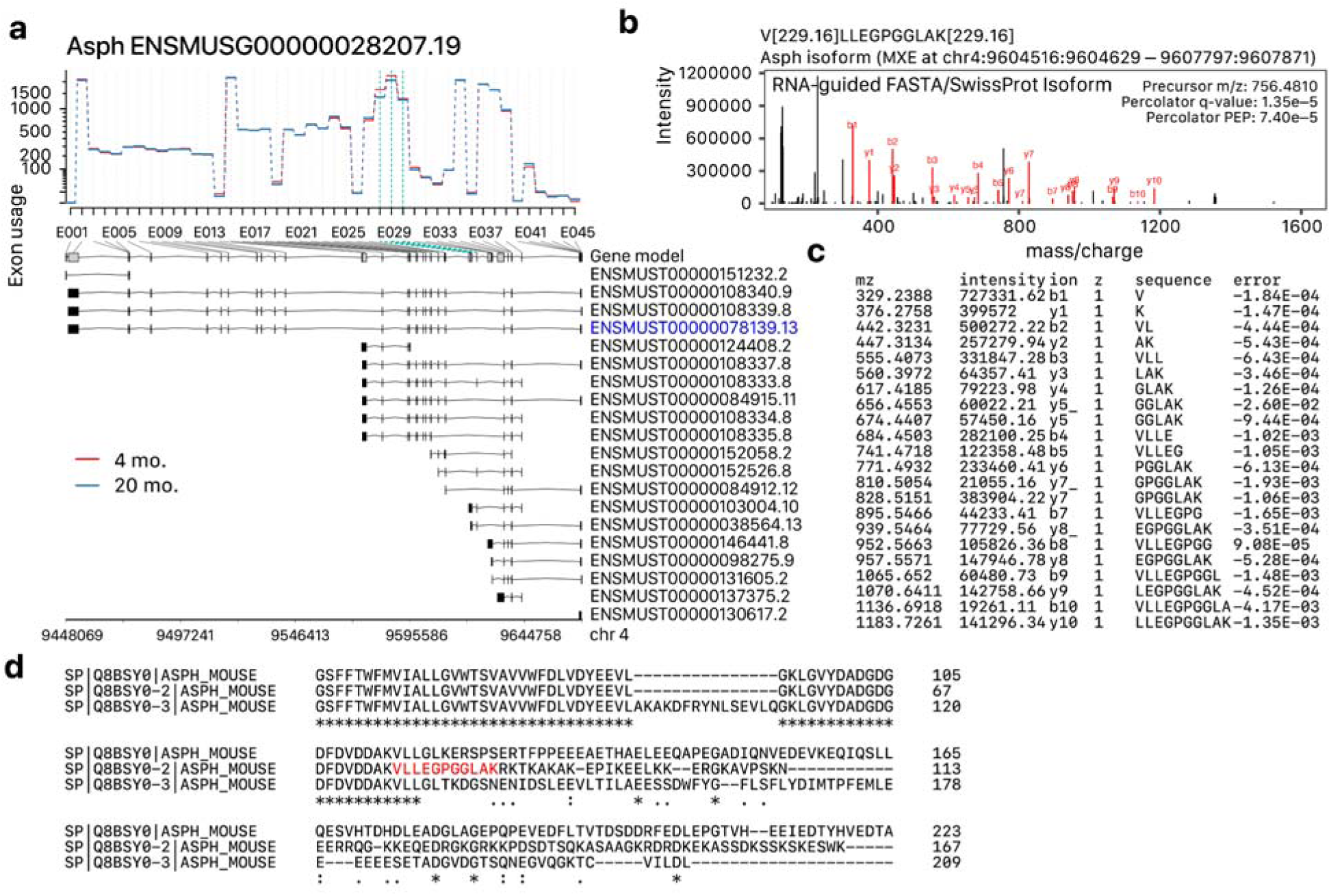
Putative identification of an alternative peptide in Asph. **a**. Exon usage map of Asph. Disjoint exonic parts E028–E030 corresponding to a region of exons not found in the canonical Asph transcript (blue) are highlighted in green. The exons in the region are found in shorter transcripts in the gene model corresponding to cardiac junctin. **b**. The best-hit peptide spectrum match for a unique peptide sequence corresponding to the junctin isoform that is identifiable using UniProt SwissProt isoform + canonical and the RNA-guided protein sequence databases. **c**. List of matched fragment b- and y-ions in the spectrum in panel b. **D**. Clustal O local sequence alignment of ASPH canonical and isoform sequences in UniProt SwissProt. The identified peptide is highlighted in red.

## Acknowledgments

This study was supported in part by NIH award F32-HL149191 to YH; University of Colorado UROP Mini-Grant to JW; NIH awards R03-OD032666 and R00-HL144829 to EL; NIH awards R00-HL127302, R01-HL141278, R21-HL150456 to ML; and University of Colorado Consortium for Fibrosis Research and Translation funding to ML.

## Notes

### Competing Interest Statement

The authors have declared no competing interest.

